# High performance sorting of motor unit action potentials with EMUsort

**DOI:** 10.64898/2026.01.06.697952

**Authors:** Sean O’Connell, Jonathan A. Michaels, Runming Wang, Sahit Mamidipaka, Manikandan Venkatesh, Nevin Aresh, Marius Pachitariu, J. Andrew Pruszynski, Samuel J. Sober, Chethan Pandarinath

## Abstract

Understanding how neural signals control muscle activity during behavior is a key challenge in motor neuroscience. To this end, recent advances in intramuscular multielectrode arrays have enabled high-quality multichannel recordings of many motor unit action potentials (MUAPs) in freely moving subjects. However, identifying individual MUAP events within multichannel recordings is a significant challenge for existing spike sorting methods, which are typically optimized for identifying action potentials from neurons in the brain. To overcome this challenge, we developed the Enhanced Motor Unit sorter (EMUsort), an extension of Kilosort4 (KS4) that achieves high-performance MUAP spike sorting. We applied EMUsort to high-resolution intramuscular recordings from rat forelimb during locomotion and monkey forelimb during a reaching task. EMUsort improves upon prior methods by addressing key challenges encountered with MUAP datasets, including: 1) long time delays across electrodes due to propagation along muscle fibers, 2) more complex waveform shapes compared to neuronal action potentials, and 3) a high degree of MUAP overlap due to cumulative motor unit recruitment. We compared EMUsort to existing spike sorting methods quantitatively using simulated datasets that closely emulated the rat and monkey datasets we recorded. EMUsort provided median error rate reductions of 67.5% and 49.9% during periods of high motor unit activation for the rat and monkey datasets, respectively. In sum, EMUsort provides a substantial improvement to MUAP spike sorter accuracy, especially during regions of high MUAP overlap, in an easy-to-use software package.

## Introduction

Motor neurons control muscle force output by sending patterns of action potentials to muscle fibers via neuromuscular junctions (Cohen & Gans, 1975; Fuglevand et al., 1993a; Marshall et al., 2022; Srivastava et al., 2017). Each motor neuron controls a set of muscle fibers in a functional unit called a “motor unit” (Heckman & Enoka, 2012). Understanding how these motor units are controlled and coordinated is therefore central to answering larger questions about behavioral control (Bräcklein et al., 2022; Hodson-Tole & Wakeling, 2009; Lemon, 2008). Neuroscientists have developed various electrophysiological methods both to record and to identify when individual motor units activate by measuring individual motor unit action potentials (MUAPs) in muscles (Farina & Gandevia, 2024; Gohel & Mehendale, 2020). These muscle recordings provide an accessible and robust measure of motor neuron activity because each action potential fired by a motor neuron triggers muscle fiber activation with nearly absolute certainty in healthy animals (Heckman & Enoka, 2012). While dense intramuscular electrode arrays provide new opportunities to record from populations of motor units (Chung et al., 2023; Metallo et al., 2011; Muceli et al., 2015, 2022; Oldroyd et al., 2025), it can be difficult to distinguish activity from individual motor units when multiple MUAPs occur on the same channels simultaneously.

In this work we introduce the Enhanced Motor Unit sorter (EMUsort), which achieves state-of-the-art performance in identifying MUAPs from dense intramuscular recordings. EMUsort builds on Kilosort4 (KS4), a high-performing spike sorter for multichannel brain recordings (Magland et al., 2020; Pachitariu et al., 2016, 2023, 2024; Steinmetz et al., 2021). EMUsort includes several innovations designed to address sorting challenges that are unique to high-density intramuscular recordings such as cross-channel time delays, complex waveform shapes, and high degrees of waveform overlap. We used Myomatrix arrays to collect high-density intramuscular recordings from both rat and monkey (triceps and biceps brachii) and use these recordings here to create realistic simulated datasets, allowing quantitative comparison of spike sorting performance in cases where ground truth spike times are known. We demonstrate that EMUsort provides a reliable improvement in spike sorting accuracy relative to alternative methods when applied to simulated rat and monkey data. These performance differences were greater during periods of higher muscle activity, which are more challenging to sort given the higher preponderance of overlapping MUAPs. Thus, EMUsort achieves state-of-the-art performance for identifying MUAPs from high-density intramuscular recordings. In addition to EMUsort’s utility for spike sorting, our method for building realistic simulated datasets provides an additional tool for future studies examining EMG signal properties in other muscles and species. All tools and datasets developed for this work are openly released for use by the community (O’Connell & Michaels, 2025; O’Connell, 2025; O’Connell & Wang, 2025).

### Background: Building on Kilosort4

To explain how EMUsort builds on KS4, we first give an overview of the basic sequence of operations performed by KS4. We then describe challenges inherent to MUAP data and how these challenges are addressed with particular components of EMUsort.

The effectiveness of EMUsort for sorting motor unit data depends on a series of modifications to KS4 that take into account the differences between electromyographic (EMG; muscle) and neural data. However, EMUsort retains much of the overall structure and many subroutines of KS4. The first processing stage of KS4 (and EMUsort) uses variance-based thresholds on each channel to extract a set of time-separated (non-overlapping) waveforms to be used as single channel templates. This set of non-overlapping waveforms is expected to capture the entire range of true spike waveform shapes present in the dataset, so their selection is a critical initialization for the downstream template matching and clustering stages of KS4. As described in Results (see “EMUsort captures the complexity of motor unit waveforms”), EMUsort modifies this stage to allow an arbitrary number of amplitude thresholds (instead of just one threshold as in KS4) for finding all non-overlapping representative waveform shapes. The use of multiple amplitude thresholds allows EMUsort to capture a larger and more varied set of MUAP waveforms for use in this first processing step.

This set of non-overlapping waveforms is used for two purposes. In KS4, these are clustered with K-means and the cluster centers are used to define the unique waveform shape for each “simple template,” which are used for the first convolutional matching stage and are defined by repeating a representative waveform shape surrounding a central channel, with spatially decaying amplitude across channels (Pachitariu et al., 2024). EMUsort takes the same approach, but the default number of clusters is increased to produce more templates for capturing the wider range of shapes usually present in MUAP datasets. Second, principal components (PCs) are computed across this set of non-overlapping waveforms for later representation of all waveforms in the dataset during subsequent matching and refinement stages. EMUsort similarly uses this low-dimensional representation of waveforms, although it increases the default number of PCs (relative to KS4, see *n_pcs* in Table 5) used to represent motor unit waveforms to reflect the greater waveform complexity of MUAPs compared to neural action potentials.

KS4 next classifies spike features using graph-based clustering methods, in which spike time and waveform shape criteria are used (based on known statistics of neural spikes) to split and merge candidate “clusters” of spikes arising from the same unit. For the time criteria, cross-correlograms are computed between spike times across clusters to minimize refractory period violations, which indicate contamination from noise or from other neurons with different cluster designations. For the template waveform shape criteria, the shapes are refined by splitting any cluster with a bimodal distribution, which is assumed to reflect the presence of more than one unit. These clustering procedures are central to Kilosort’s effectiveness and are deployed similarly in EMUsort, although EMUsort provides additional modifications that account for key differences between neural and muscular data, as described below.

### Background: Motor unit datasets present unique spike sorting challenges

Motor unit waveforms recorded with intramuscular arrays have several characteristics that make sorting their waveforms more challenging than sorting spikes recorded from neurons. First, MUAPs generally have more complex waveform shapes than cortical neuron action potentials, reflecting the contributions of many muscle fibers distributed widely throughout the muscle (Figure 2A). In contrast, spike waveforms from neurons arise from a single cell with a soma that is typically much smaller than any of the individual fibers that comprise a motor unit (Buzsáki et al., 2012; Gold et al., 2006). Second, because depolarization takes time to travel along the muscle fibers, we generally observe larger time delays between voltage peaks on neighboring EMG channels than the near-zero delays between somatic spikes obtained during dense multisite recording of neurons (Figure 1C, F). Finally, more motor units are recruited rapidly as muscle force increases, in contrast to cortical neurons, so that MUAPs recruited early are obscured by MUAPs recruited later as force increases, making the MUAPs much more difficult to sort at higher activation levels (Figure 1B, E). Below we detail each of these challenges.

**Figure 1.**
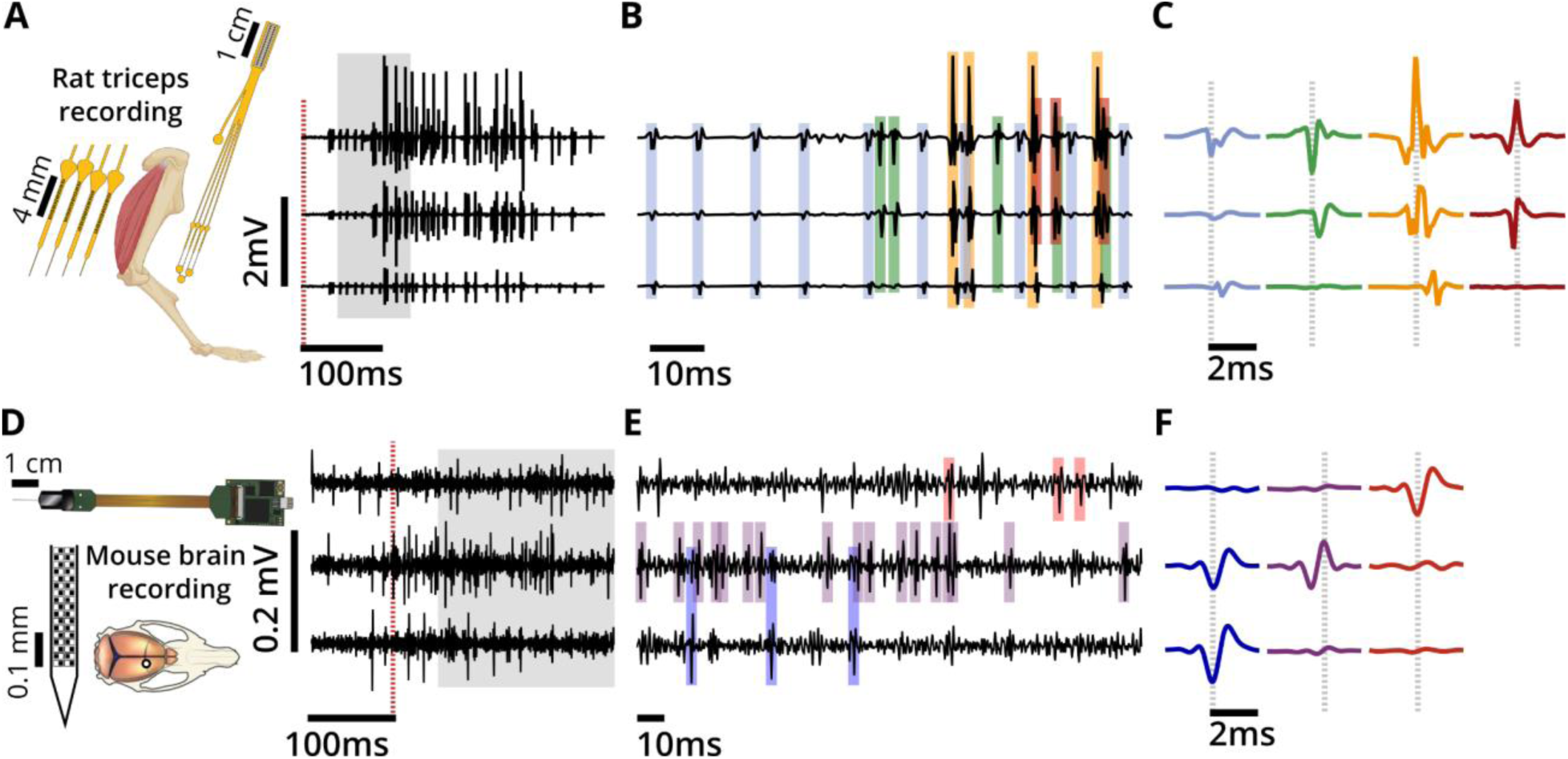
Motor units exhibit more complex action potential waveforms than neurons. (a) Example recording with a Myomatrix array in a rat triceps brachii (lateral head) during the stance phase of a single stride cycle during treadmill locomotion. The red dashed line indicates the beginning of stance. Channels were spaced in the muscle by 1 mm. (b) Grey region expanded from panel (a), with 4 unique MUAP shapes identified by colored highlights. (c) Extracted mean waveforms from each of the identified MUAPs in this recording. Grey dashed lines illustrate temporal delays across recording channels. (d) Neuropixels 1.0 recording from primary motor cortex layer 5 in a mouse during a decision-making task. The red dashed line indicates movement onset. Channels were spaced in the brain by 40 and 120 μm. Data and resources for this panel are from (International Brain Laboratory et al., 2024; Kristensen & Jörntell, 2023). (e) Grey region expanded from (d), with 3 unique neural waveforms identified by colored highlights. (f) Extracted mean waveforms from this recording. Note the significant conduction delay across channels in motor unit data (c) but not neural data (f) as well as the greater waveform complexity of MUAPs as compared to neural action potential waveforms.

**Figure 2.**
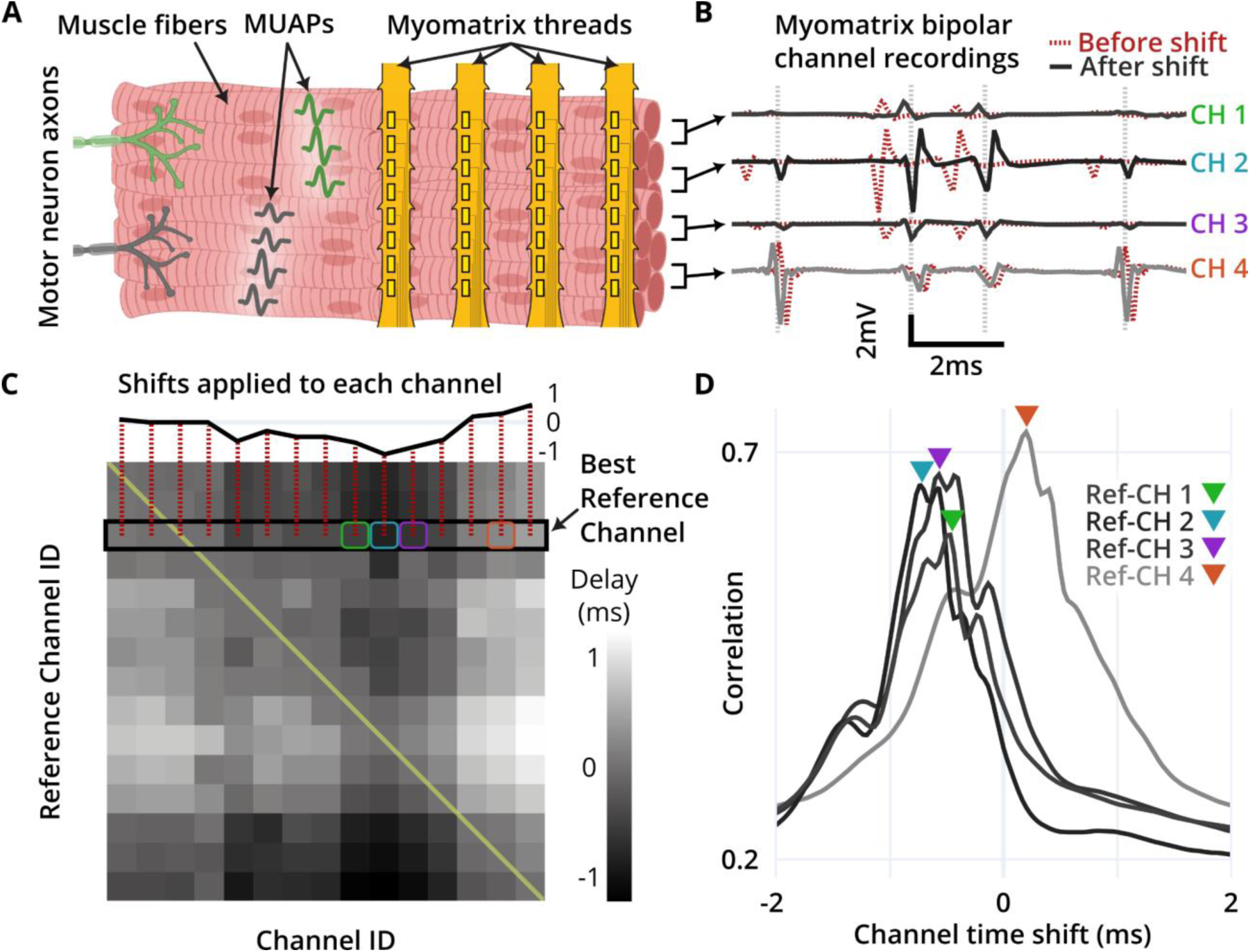
Electrode placement within a muscle determines relative time delays of cross-channel signals for a given MUAP. By comparing cross-channel correlations across a range of delays, motor unit action potential (MUAP) time alignment can be performed to improve spike sorting. (a) Schematic depicting Myomatrix electrode array and MUAPs propagating along muscle fibers. (b) EMG data from a rat triceps during treadmill locomotion. Four channels from the original recording are shown by red dashed lines. The black lines show the data after the channels have been shifted. (c) Grayscale map of temporal delays that maximize channel correlations for each reference channel. For each row, unshifted reference channels lie along the yellow diagonal. The shading in each square indicates the best channel shift to apply to maximize the correlation with the reference channel, with lighter shades representing higher correlations. The black rectangle shows the best choice of reference channel and shifts that produced the highest sum of correlations. Colored squares show the true position of channels taken from the full recording for use in this figure. (d) Correlation values for each of the 4 selected channels. Peak correlations are marked by colored triangles, indicating the channel time shift that maximized the correlation with the reference channel.

#### MUAP waveforms can be time-delayed across channels

An additional challenge in motor unit spike sorting is the delay between the arrival times of a MUAP at spatially separated electrode contacts (Figure 1C). This phenomenon does not occur as markedly because of the much smaller size of neural soma compared to muscle fibers and the much closer packing of, for example, Neuropixels contacts compared to the inter-contact spacing of Myomatrix arrays (Buzsáki et al., 2012; Chung et al., 2023; Steinmetz et al., 2021). As a result, inter-channel delays are far larger in EMG recordings compared to neural recordings (Merletti & Farina, 2008; Swadlow & Waxman, 2012; Thompson et al., 2018). With potentially substantial time delays across channels in recordings of MUAPs (Figure 1C), the spike sorting performance is degraded due to the increased capacity required to be able to reconstruct the shifted waveform shapes with a shared temporal basis. This is especially true when the temporal basis is constructed from centrally-aligned single channel waveforms, as in KS4.

#### MUAP waveforms have more complex shapes than action potentials in the brain

The action potential of a single motor unit typically has a more complex waveform than the action potential of a neuron (Figure 1C, F), reflecting the fact that each motor unit waveform results from the summation of voltage deflections from each of the many muscle fibers innervated by a single motor neuron. Moreover, each EMG electrode has a different location relative to the set of activated muscle fibers, resulting in different waveform shapes across channels. Similarly, the polarity and number of phases in the MUAP are determined by the electrode position relative to the neuromuscular junction on each fiber and slightly different conduction speeds caused by slight differences in the size of each fiber (Konstantin et al., 2020). All these influences contribute to the greater waveform complexity of MUAPs, with multiple phases and large positive peaks. In contrast, waveforms recorded from cortical neurons are less variable across neurons and are often biphasic with a larger negative peak occurring before a smaller positive peak. All these factors tend to produce simpler spike waveforms, even across different cortical areas (Buzsáki et al., 2012; Gold et al., 2006; Quian Quiroga & Panzeri, 2013; Someck et al., 2023). As quantified below, the greater waveform shape complexity of MUAPs results in high error rates when KS4 is applied to motor unit data (without the appropriate modifications presented here).

#### MUAP waveforms tend to overlap frequently during forceful behavior

Differences between the firing dynamics of motor units and neurons also create unique challenges for MUAP spike sorting. First, whereas cortical neurons fire spontaneously even in the absence of body movement, many motor units only fire during active movement with higher levels of force. Second, whereas earlier-recruited (lower force) motor units have MUAPs with smaller amplitudes, MUAPs from later-recruited (higher force) motor units are generally much larger in amplitude and time width. These factors combine to produce more concentrated MUAP activity, with the smaller MUAPs becoming much harder to detect in the presence of larger MUAPs. The result is much greater action potential overlap rates in high-resolution EMG datasets compared to neural recordings. To maintain high spike sorting performance with MUAP datasets, this increased rate of waveform overlap must be accounted for, to be able to further our understanding of the neural control of motor behavior. We therefore identified and addressed failure modes in MUAP identification caused by these bursts of activity (e.g., using multiple amplitude thresholds, outlier rejection, and cross-channel alignment) that improved the performance of deconvolution procedures used in KS4 (Pachitariu et al., 2024). The developments we implemented in EMUsort were therefore essential to better account for the biological differences between neurons and motor units.

## Results

The organization and design of EMUsort provides usability and runtime benefits. EMUsort provides a command line tool with an emphasis on straightforward configuration and producing fast results for the user. To do this, we built the command line interface with Python and leveraged the open-source SpikeInterface library (Buccino et al., 2020) for data loading, channel map creation, preprocessing, bad channel rejection, sorter job parallelization, and quality metric calculation. EMUsort removes the need to write code to perform many individual tasks: loading common dataset types, adjusting filter parameters, sorting specific time intervals of individual recordings, concatenating multiple recordings, specifying channel groups to be sorted separately in series, and running parameter “sweeps” across sorting parameter values in parallel. EMUsort also computes a composite score to estimate overall sort quality (see “Producing Composite Scores for Agnostic Estimation of Sort Quality” in Methods), which is appended to the name of each sort output folder to indicate to the user which sorts are likely better than others. To our knowledge EMUsort is the first spike sorting tool that enables the user to perform parameter sweeps without writing any code outside of the configuration file and provides automatic performance estimates (composite scores) for each sorting run.

The configuration and execution of EMUsort are designed to optimize user convenience and fast runtime. Most common operations in EMUsort can be controlled from the primary configuration file (*emu_config.yaml*), which can be launched directly in the terminal for fast verification or adjustment before a spike sorting run, or to set up multiple runs in parallel. The user configures EMUsort to sweep combinations of selected parameters by editing *emu_config.yaml* (which by default sweeps the parameters *Th_single_ch*, *Th_universal*, and *Th_learned*; see Table 5). EMUsort then distributes the number of jobs evenly across the selected GPUs, and when all sort jobs are complete, it handles all data writing operations asynchronously to minimize overall run time. To quantify EMUsort runtimes and compare these with the time required to execute another state-of-the-art motor unit spike sorter, when we ran a parameter sweep in parallel across 25 parameter combinations for a 10 minute simulated rat dataset with 8 channels and 10 motor units, all 25 sort runs and asynchronous data writes finished in 10.9 minutes (effectively 26.1 seconds per sort). In contrast, when we swept the same parameters sequentially, it took 50.7 minutes to complete (121.7 seconds per sort, on average). These run times are advantageous in comparison to alternative available MUAP sorting softwares, such as runs with MUedit (Avrillon et al., 2024), in which each individual run on the same dataset, using a single parameter set, took greater than 4 hours. With a similar parameter sweep using a 10 minute simulated monkey dataset with 16 channels and 5 motor units, we found an even greater difference. EMUsort took 8.5 minutes to complete 25 sort jobs in parallel (effectively 20.4 seconds per sort) whereas each individual MUedit sort took greater than 31.6 hours. EMUsort therefore combines a user-friendly command interface with comparatively rapid data processing.

### EMUsort extends Kilosort4 to address spike sorting challenges with MUAP recordings

To achieve high-performance spike sorting of MUAP data, EMUsort extends KS4 to correct for the time delays present across channels, account for increased waveform shape complexity, and ensure the identification of both earlier- and later-recruited motor units. Below, we describe our main modifications to the KS4 codebase.

#### EMUsort corrects for time delays across channels to improve MUAP representation

Because MUAPs take time to propagate along muscle fibers, electrode placement greatly affects the time delays measured across channels (Figure 2A). When we observed time delays in our EMG data, the algorithm of KS4 tended to produce multiple clusters for the same underlying MUAP pattern due to improper “splitting” of delayed versions of the same MUAP. This occurred more frequently when cross-channel delays were greater than half the template window, where parts of the overall MUAP pattern can drop out of the template window, reducing the information available of the true waveform identification during template matching. Even smaller delays across channels can create difficulty in representation, because the largest peak of the temporal components align to the center of the window.

To reduce the negative impact of cross-channel time delays on spike sorting performance, EMUsort quantifies and applies time shifts to maximize the time alignment of multichannel MUAPs (Figure 2B). In high-SNR recordings, cross-channel correlation serves as a proxy for MUAP alignment, so EMUsort computes the cross-correlation matrix between each channel’s unshifted data and another channel’s data that has been time-shifted between -2 ms to +2 ms (e.g., -60 to +60 samples when using a 30 kHz sampling rate). The result is a 3-dimensional matrix of correlations between all channel pairs, for all lags. We find the indexes of the maximum correlation values along the channel shift dimension to yield the “best” shift amounts for each channel pair (Figure 2D). The entire 2-dimensional matrix of best shifts is shown in (Figure 2C). EMUsort uses the row that produces the highest sum of corresponding correlations across channels to determine the best “reference” channel and the shifts to be applied to the other channels. This delay-removal procedure can be enabled or disabled with the *remove_chan_delays* parameter (see Table 5).

#### EMUsort captures the complexity of motor unit waveforms

KS4 uses spatial components (also called channel weights) and temporal components in combination to represent the multichannel waveforms for each MUAP. The temporal components are formed from the PCs of the non-overlapping waveforms (described in Background, see “Kilosort4 spike sorting details” in Methods for more detail), and the channel weights define the magnitude of each temporal component to best represent the waveform shape on each channel during template matching (Figure 3A). In this depiction, the temporal components are shared across all waveform shapes, but a different *U* matrix would be used for each MUAP and each channel.

**Figure 3.**
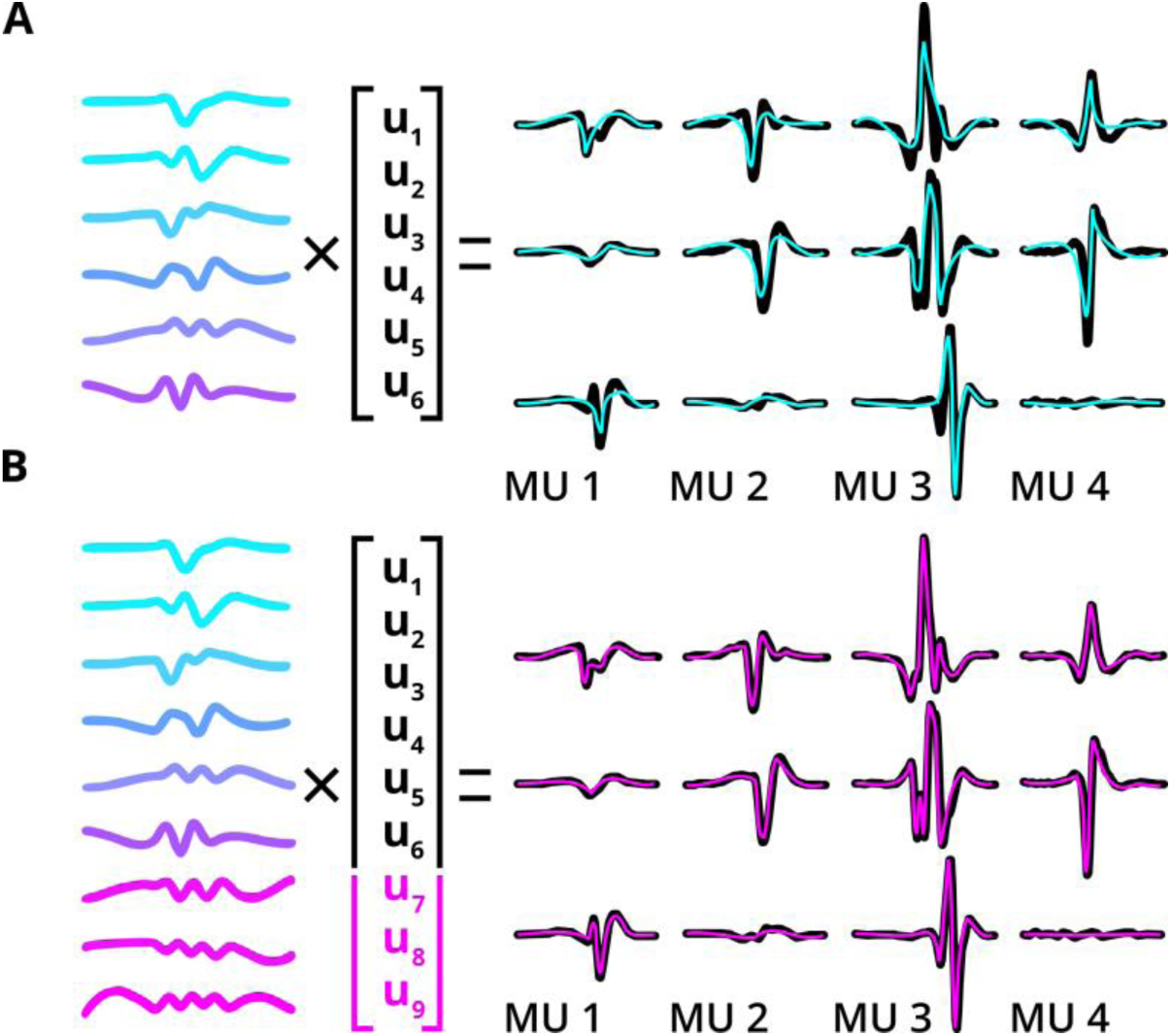
Increasing the number of temporal components provides the capacity to reconstruct a wider range of complex MUAPs across channels. Temporal components are shown on the left, which are effectively combined during template matching to represent all spike shapes in the dataset. The shape for each motor unit and channel uses the same temporal components but different values for *U*. Additional temporal components are shown in magenta. (a) Example mismatch between the true multiphasic spike shapes and the reconstruction when using only 6 temporal components with MUAP data. (b) Example of better fitted MUAP spike shapes after increasing the number of temporal components from 6 to 9.

To allow EMUsort to capture the higher complexity of MUAP waveforms, we increased the number of temporal components (controlled by *n_pcs*; see Table 5) to better represent the statistics of MUAPs. Increasing the number of components provides larger capacity to represent a wider range of MUAP shapes. Using more temporal components also allows EMUsort to represent higher frequency components of MUAPs, allowing higher precision during later convolutional processing stages. We found that increasing the number of temporal components enabled more faithful reconstruction of the original MUAPs (Figure 3B). Sorting performance improved when increasing the number of components from 6 (KS4 default) to 9 but did not improve for larger increases in component number. Finally, we also observed some MUAP patterns being missed altogether, i.e., with no final cluster being assigned to those patterns. We determined that within the set of non-overlapping threshold crossings, some clusters were not assigned to any simple template. This was largely addressed by increasing the number of simple templates (also changed from 6 to 9 between KS4 and EMUsort; controlled by *n_templates*, see Table 5), which allowed the simple templates to better span the full range of waveform shapes across all channels in the dataset.

#### Cleaner “simple template” initialization with EMUsort enhances all downstream sorting stages

We modified the template initialization of KS4 in two additional ways to improve the quality of MUAP representation by the simple templates. (Again, the simple templates perform convolutional matching to initialize the spike classifications and are created by repeating a waveform from the set of non-overlapping spikes across multiple channels at decaying amplitudes around a central channel.) First we implemented the ability to use multiple single channel thresholds and aggregate the non-overlapping spikes across thresholds to ensure capturing all unique MUAP waveforms present in the dataset. Second, we introduced a new outlier rejection method to prune inconsistent MUAP shapes from the set of non-overlapping crossings prior to creating the simple templates.

As described in the Introduction and shown in (Figure 4A-B), KS4 uses voltage threshold crossings (specifically those that define waveforms that are non-overlapping in time with other waveforms) to identify spike waveforms to use for creation of the simple templates. In EMG data, the earlier-recruited motor units (MU1 and MU2 in Figure 4A-D) are relatively easy to isolate using a single threshold value, since only one or a very few motor units are active. In contrast, MUAPs from motor units recruited later (MU3 in Figure 4A-D) only appear after many other motor units have been activated. As a result, we found that when using only a single amplitude threshold (as in KS4), MUAPs from later-recruited motor units were often excluded from the initial set of simple templates (just as MU3 is excluded in Figure 4A-B), leading to that unit being omitted from the final sort output. We addressed this problem in EMUsort by allowing the user to select multiple thresholds (e.g., both T_1_ and T_2_ in Figure 4; this is controlled by setting *Th_single_ch* as a list with multiple values; see Table 5) to facilitate better representation of all unique MUAP shapes present in the dataset (Figure 4E), even those that tend to only appear in regions of high activity.

**Figure 4.**
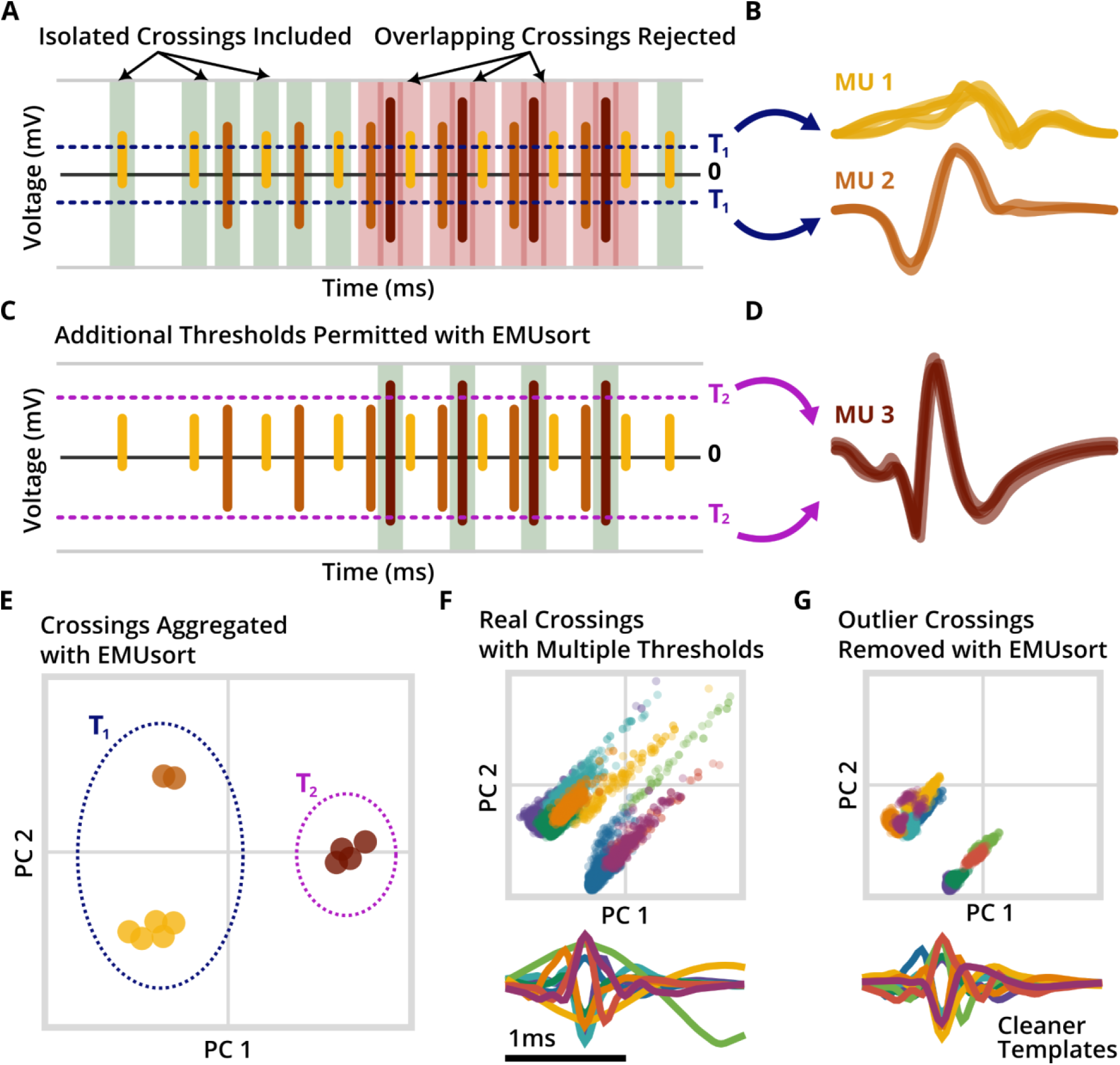
EMUsort provides improved methods for template initialization. (a) Time series graph showing the default procedure KS4 uses for finding non-overlapping spike examples to use as data for creating “simple templates” (see main text). The non-overlapping crossings (green highlights) are included in formation of the simple templates, but the overlapping crossings are rejected. (b) Depiction showing the use of a single threshold (as in KS4) to extract waveforms. The blue threshold, T_1_, extracts several examples of MU 1 and 2 (colored yellow and orange), for use with template creation, but none of MU 3 due to the isolation criteria. (c, d) Using additional thresholds (T_2_; new in EMUsort) captures waveforms of various amplitudes (e.g. larger, more crowded MUAPs like MU 3, colored brown). (e) PCA projections of the 61-point waveforms selected by using all thresholds combined, with the same color coding. (f) All PCA projected crossings from a real rat triceps brachii recording during treadmill locomotion (PCA computed using all isolated waveform shapes from the recording). Using K-means, the cluster centers are identified and used for simple template shapes. The bottom plot shows the 61-point waveforms corresponding to each cluster center. (g) Outlier removal in EMUsort with the same biological data. Low density, outlier spikes are removed with HDBSCAN, and the remaining spikes are clustered as normal with K-means., where outlier removal has resulted in a cleaner set of simple template shapes.

EMUsort also includes novel features to reduce the impact of “outlier” waveforms in EMG data. We found the spike detection method would sometimes mistakenly include movement artifacts (rare but large voltage transients resulting from electrode movement rather than muscle activity) in the set of non-overlapping crossings, which would contaminate the clustering process and warp some of the simple template shapes because they are formed from the centers of K-means clusters (Figure 4F). To avoid this contamination issue, EMUsort employs Hierarchical Density-based Spatial Clustering of Applications with Noise (HDBSCAN) to model the spike waveform clusters in the full feature space (61 samples by default) based on local densities of points, and perform outlier rejection by excluding spike waveforms that do not fall within defined clusters (Figure 4G) (Campello et al., 2013, 2015). After the outlier waveforms are removed from consideration, the reduced set of non-overlapping crossings are then clustered with K-means, with the cluster centers taken to define the simple templates.

### EMUsort improves sorting in biological and simulated datasets

To assess EMUsort performance when applied to EMG data, we collected data from rats and monkeys during behavioral tasks (see Methods) and performed many spike sorting runs using EMUsort and default KS4. Inspection of the sorted output suggested lower type I error with EMUsort (i.e., fewer false positives), as seen by reduced refractory period violations in the autocorrelogram. There were also cases where EMUsort labeled unique motor units that were missed by KS4, indicating lower type II error (i.e., fewer false negatives). EMUsort appeared to perform particularly well during bursts of high muscle activity when overlaps of many different motor unit waveforms were more likely to occur (Figure 5A-B), highlighting the need to quantify sorter performances across different levels of muscle contraction.

**Figure 5:**
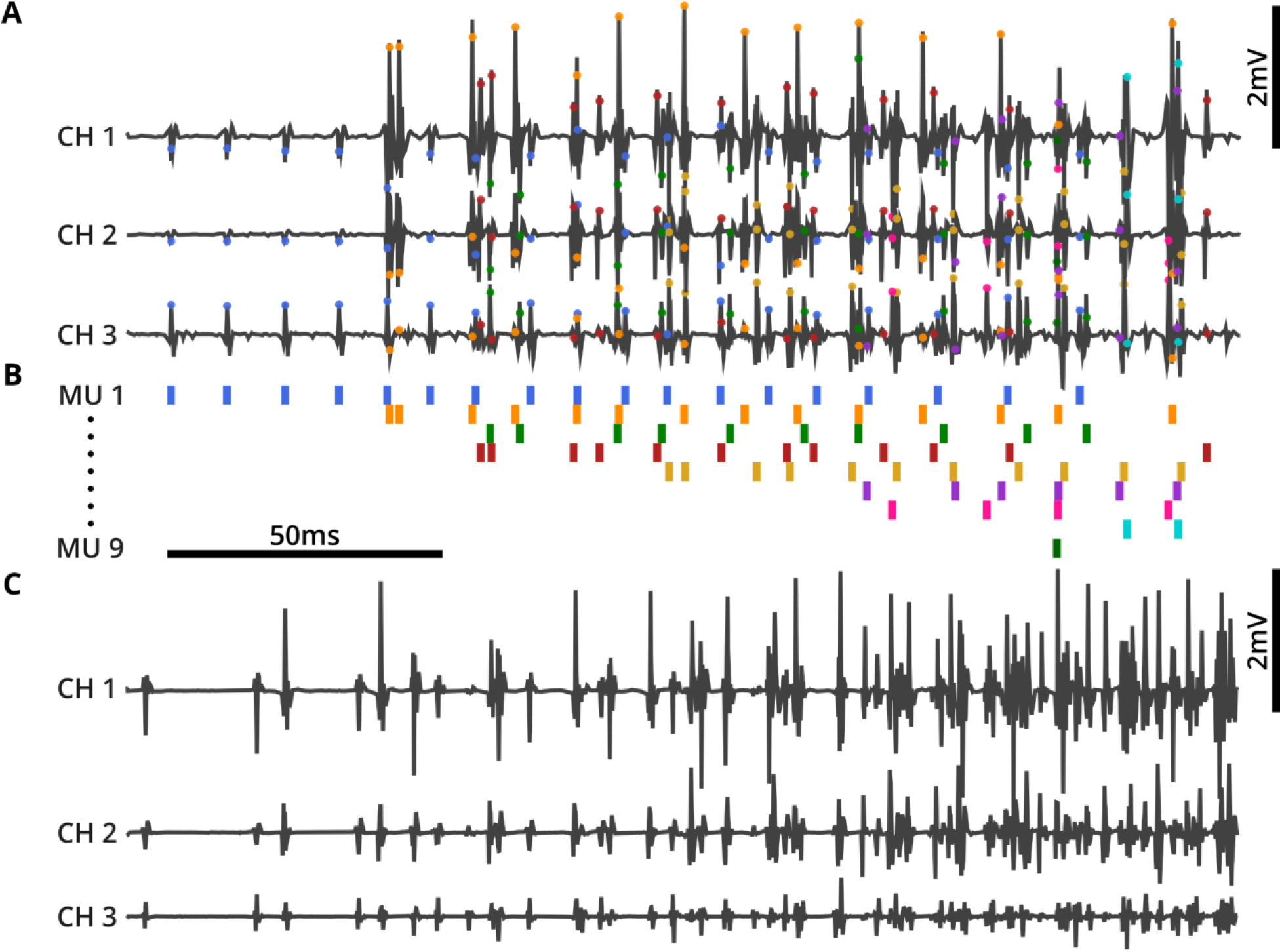
Qualitative sorter performance comparison of biological data and a representative example of simulated data. Multichannel MUAP spike sorting can be difficult to validate by visual inspection, necessitating quantitative methods for performance assessment. Here we show the sorting of biological data and a sample of the multichannel simulation created from the real dataset. (a) Real EMG data recorded from triceps brachii in a rat during treadmill locomotion. (b) Event plot showing the visually “best” spike sorting result from EMUsort. (c) Example of simulated EMG “data” using simulated spike times (see text).

To quantify the performance of each sorter, we used the waveform shapes and interspike intervals from our biological EMG recordings in rats (Figure 5A) and monkeys to create simulated EMG datasets (Figure 5C shows a rat simulation example). This approach provided simulated voltage data with known ground truth spike times for 10 simulated units from the rat data and 5 simulated motor units from the monkey data with species-appropriate MUAP shapes. For both simulations, the shapes and amplitudes of each unique MUAP were distributed across different channels as in the empirical data. For the rat simulation the 10 MUAPs were simulated across 8 channels, and with monkey simulation, the 5 MUAPs were distributed across 16 channels (see “Generation of simulated MUAP datasets” in Methods). We used these simulated datasets to run parameter sweeps and conduct a quantitative comparison of sorting performance between EMUsort and KS4. We also ran the same performance comparisons with MUedit (Avrillon et al., 2024), across a range of parameter settings, and consulted with the MUedit developers to validate our parameter choices. For information about parameters used, see (Table 3) in Methods.

#### EMUsort achieves substantially higher sort accuracy on a simulated rat dataset

EMUsort consistently outperformed KS4, reflecting the many changes we made to KS4 to account for the different statistics of motor unit versus neural activity. We conducted 25 sort runs each with EMUsort and KS4, and 4 sort runs with MUedit, using a rat-based simulated dataset with 10 motor units firing across 8 channels with a duration of 10 minutes (see “Generation of simulated MUAP datasets” in Methods). Evaluating parameter sweeps showed EMUsort produced the best distribution of mean accuracies across the simulated units. The sorter accuracy statistics are shown in (Table 1), where each point in the distributions is a mean across unit accuracies within each sort. We next quantified performance in the face of MUAP overlaps, which are particularly challenging and highly prevalent in intramuscular recordings during periods of high force output. We defined overlaps as events where neighboring MUAP times differed by half the template window or less (1 ms and 2 ms for the rat and monkey datasets respectively). The resulting accuracy statistics of these comparisons for all sorts with the rat data simulation are shown in (Table 1 and Figure 6). Alternative measures of accuracy and error rates used in (Pachitariu et al., 2016) were computed separately and are shown in (Table 1 - table supplements 1-2). We found that not only does EMUsort have a higher median and grand mean accuracy across sort runs when compared to KS4 and MUedit, but the performance gap is substantially larger in the regions of data with higher overlap (Figure 6A-B). EMUsort may therefore be particularly valuable for scientific investigations that focus on dynamic, unrestrained movements or tasks with high force outputs when overlaps are much more likely to occur.

**Figure 6:**
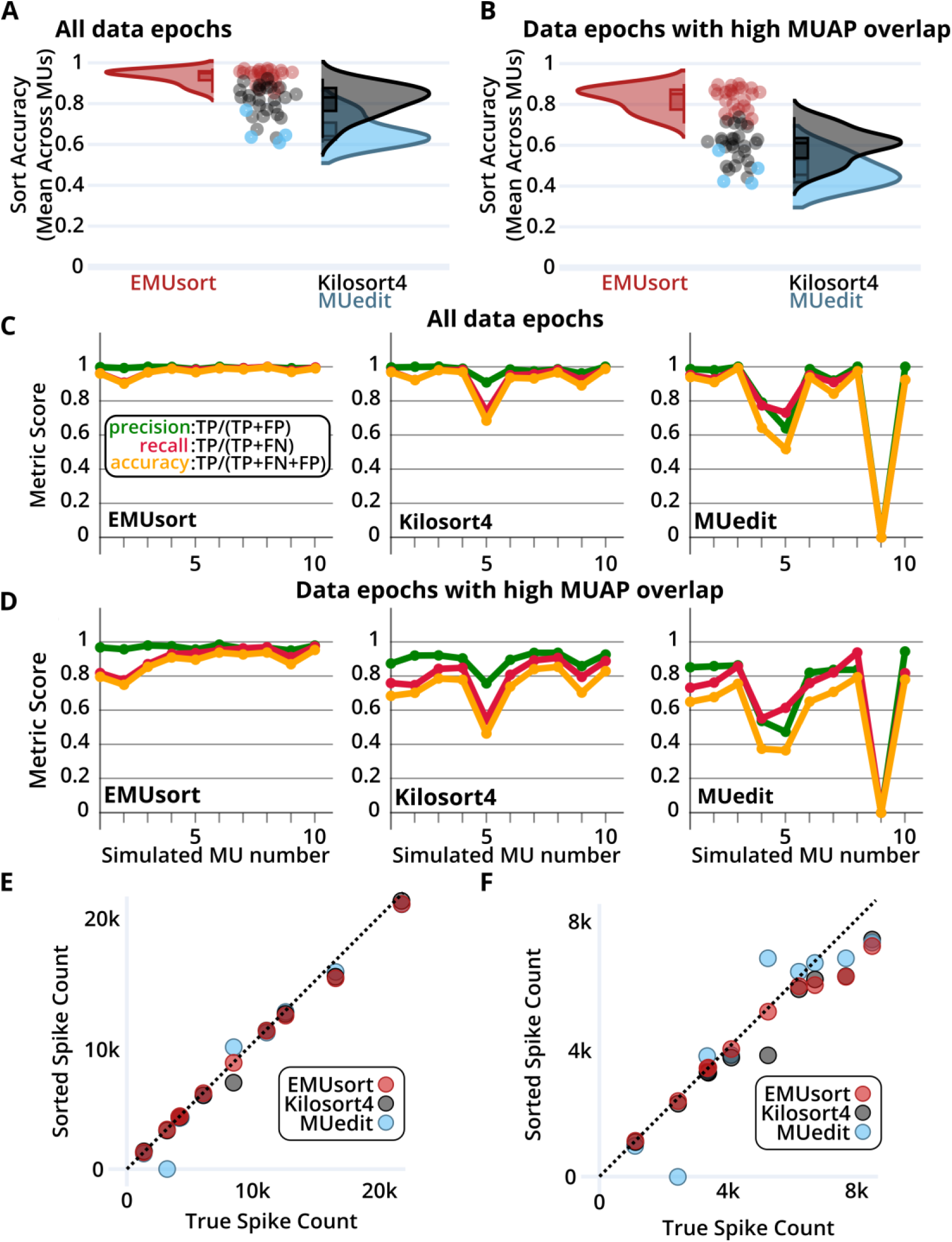
EMUsort achieves high performance with rat-based simulations using real MUAP recordings from triceps brachii during locomotion. (a) Sort accuracy for EMUsort and KS4 across a parameter sweep with 25 combinations. Sort accuracies from four MUedit runs are also shown. Each point shows the mean accuracy across ground truth units for each sort run. (b) Same metrics as (a) but the accuracy calculations were computed only data epochs that included highly overlapped spikes, i.e., only MUAPs within at least 2 ms of another MUAP. This assesses how each sorter performs in the most challenging examples in the simulated dataset. In (a, b), the distribution plot is created with kernel density estimation for visualization. (c) Per-unit performance metrics from the best sort for each sorter using all spikes in the rat dataset, arranged from highest to lowest spike count. (d) Per-unit metrics for the same best sorts as in panel (c) but computed only on overlapping MUAPs as in panel (b). (e) Scatter plot showing how well the best sort from each sorter matches the true spike counts, where paired counts for each ground truth unit are shown as a point. The dotted line indicates where the sorted count should match the true count. Note that panels (e, f) reflect total spike counts regardless of sorting accuracy. (f) Same metrics as in (e) but applied only to data with overlapping MUAPs.

**Table 1.** Sorter accuracy statistics across multiple spike sorting runs with a rat-based simulation. Runs were performed across a range of different parameters and the accuracies were computed using all MUAPs. The bracketed values correspond to results using only the overlapping MUAPs. Data points in these distributions are mean accuracies across units in each sort.

|  |  | Descriptive Statistics for Distributions of Cross-Unit Mean Accuracies |  |  |
| --- | --- | --- | --- | --- |
| Sorter | Run Count | Median (%)<br>all MUAPs<br>[overlapping MUAPs] | Mean (%)<br>all MUAPs<br>[overlapping MUAPs] | Standard Deviation (%)<br>all MUAPs<br>[overlapping MUAPs] |
| EMUsort | 25 | 94.9 [85.1] | 93.4 [82.7] | 3.4 [5.9] |
| Kilosort4 | 25 | 83.0 [61.1] | 81.5 [60.0] | 7.4 [8.0] |
| MUedit | 4 | 64.1 [45.5] | 66.5 [47.5] | 7.0 [7.4] |

We next examined performance metrics for individual motor units from the highest performing sort from each sorter, and found that for each motor unit, EMUsort consistently performed better than or approximately equal to both KS4 and MUedit (Figure 6C). EMUsort particularly outperformed the other sorters for units with lower spike counts (the units presented in (Figure 6C-D) are in descending order of spike count). This property of capturing the rarer units is likely due to incorporating multiple non-overlapping threshold crossings and including more simple templates, which allow capturing a wider range of shapes during the initial data convolutions. Further, when examining individual unit results for overlapping spikes (Figure 6D), we found that EMUsort scored the highest accuracy for all 10 simulated motor units.

Finally, we examined how well each sorter captured the total number of spikes. When including all spikes (Figure 6E), EMUsort more accurately produced the true spike count relative to other sorters. All sorters had some difficulty capturing the true spike counts during regions of high overlap (Figure 6F), especially for the higher spike count units, which tended to have lower amplitudes and could be obscured by the larger signals of simultaneously occurring MUAPs. Although MUedit appears to reflect the true spike counts for some of the higher count units (blue dots lying on the dotted line), this in fact reflects lower precision (i.e., inclusion of extraneous spikes, which compensated for missed true spikes). Detailed per-unit statistics for the best sorts of each sorter are presented in (Table 1 - table supplements 3-8).

#### Performance metrics with a simulated monkey dataset show the highest performance with EMUsort

To test whether EMUsort’s improved performance generalized beyond rodent data, we repeated the performance comparison on a separate simulated dataset created from monkey biceps brachii recordings (see Methods). We again performed a parameter sweep with 25 sort runs for EMUsort and KS4, and 4 sort runs with MUedit (see Table 4 for parameters used). Notably, the monkey dataset contained fewer unique MUAPs (5 motor units firing across 16 channels), making it less challenging to sort. As shown in (Table 2), when we included all spikes in the analysis, EMUsort achieved the best grand mean and median across all mean sort run accuracies (both greater than 97% accurate), but KS4 showed a slightly lower standard deviation. When we focused the analysis only on the overlapping spikes (with a 2ms overlap window to account for wider waveforms), the performance difference between sorters became more apparent, with EMUsort showing an 8-12% boost in accuracy during overlap events. The different performance across species is likely explained by the larger number of motor units used in the rat simulation (10 versus 5), simulated at higher firing rates, together leading to more overlaps and more opportunities for misidentification.

**Table 2.** Sorter accuracy statistics across multiple spike sorting runs with a monkey-based simulation. Runs were performed across a range of different parameters, and the accuracies were computed using all MUAPs. The bracketed values correspond to results using only the overlapping MUAPs. Data points in these distributions are mean accuracies across units in each sort.

|  |  | Descriptive Statistics for Distributions of Cross-Unit Mean Accuracies |  |  |
| --- | --- | --- | --- | --- |
| Sorter | Run Count | Median (%)<br>all MUAPs<br>[overlapping MUAPs] | Mean (%)<br>all MUAPs<br>[overlapping MUAPs] | Standard Deviation (%)<br>all MUAPs<br>[overlapping MUAPs] |
| EMUSort | 25 | 98.2 [83.9] | 97.2 [80.0] | 2.6 [9.6] |
| Kilosort4 | 25 | 95.9 [71.1] | 95.8 [72.3] | 2.3 [5.4] |
| MUedit | 4 | 93.4 [65.0] | 91.3 [65.4] | 5.1 [9.4] |

**Table 3.** Parameter sweep values for each sorter with the rat simulated dataset. With EMUsort and KS4, each pair of *Th_universal* and *Th_learned* values were used with each set of *Th_single_ch* values for a total of 25 parameter combinations. For explanations of *Th_universal*, *Th_learned*, and *Th_single_ch*, see “Kilosort4 spike sorting details”. With MUedit, each Initialization setting was used with each Contrast Function setting listed in the table, totaling 4 parameter combinations.

| EMUsort |  | Kilosort4 |  | MUedit |  |
| --- | --- | --- | --- | --- | --- |
| *[Th_universal, Th_learned]* | *[Th_single_ch]* | *[Th_universal, Th_learned]* | *[Th_single_ch]* | Initialization | Contrast Function |
| [9, 8] | [6, 9, 12, 15] | [9, 8] | [15] | Random | Kurtosis |
| [10, 4] | [6, 9, 12] | [10, 4] | [12] |  |  |
| [7, 3] | [3, 6, 9] | [7, 3] | [3] | EMG Max | Skew |
| [5, 2] | [6, 9] | [5, 2] | [9] |  |  |
| [2, 1] | [6] | [2, 1] | [6] |  |  |

**Table 4.** Parameter sweep values for each sorter with the monkey simulated dataset. With EMUsort and KS4, each pair of *Th_universal* and *Th_learned* values were used with each set of *Th_single_ch* values for a total of 25 parameter combinations. For explanations of *Th_universal*, *Th_learned*, and *Th_single_ch*, see “Kilosort4 spike sorting details”. With MUedit, each Initialization setting was used with each Contrast Function setting listed in the table, totaling 4 parameter combinations.

| EMUsort |  | Kilosort4 |  | MUedit |  |
| --- | --- | --- | --- | --- | --- |
| $[Th_{universal}, Th_{learned}]$ | $[Th_{single\_ch}]$ | $[Th_{universal}, Th_{learned}]$ | $[Th_{single\_ch}]$ | Initialization | Contrast Function |
| [9, 8] | [6, 9, 12, 15] | [9, 8] | [15] | Random | Kurtosis |
| [10, 9] | [6, 9, 12] | [10, 9] | [12] |  |  |
| [7, 6] | [3, 6, 9] | [7, 6] | [3] | EMG Max | Skew |
| [5, 4] | [6, 9] | [5, 4] | [9] |  |  |
| [12, 11] | [3, 6] | [12, 11] | [6] |  |  |

In the simulated monkey dataset, we again see higher median and grand mean accuracies with EMUsort when including all spikes in the analyses (Figure 7A). Compared to the rat simulation, the smaller gap between sorters seems to be driven by the monkey dataset being less challenging to sort (having fewer motor units than the rat simulation), so that all sorters are approaching the performance ceiling. However, as in the rat results shown in (Figure 6A-B), in the analysis of simulated monkey data the differences in performance became larger for overlapping spikes (Figure 7B), with larger separations between the medians (see bracketed results in Table 2), again with EMUsort performing best. This supports the previous results with the rat dataset that EMUsort provides enhanced MUAP sorting accuracy during data segments with high muscle activity.

**Figure 7:**
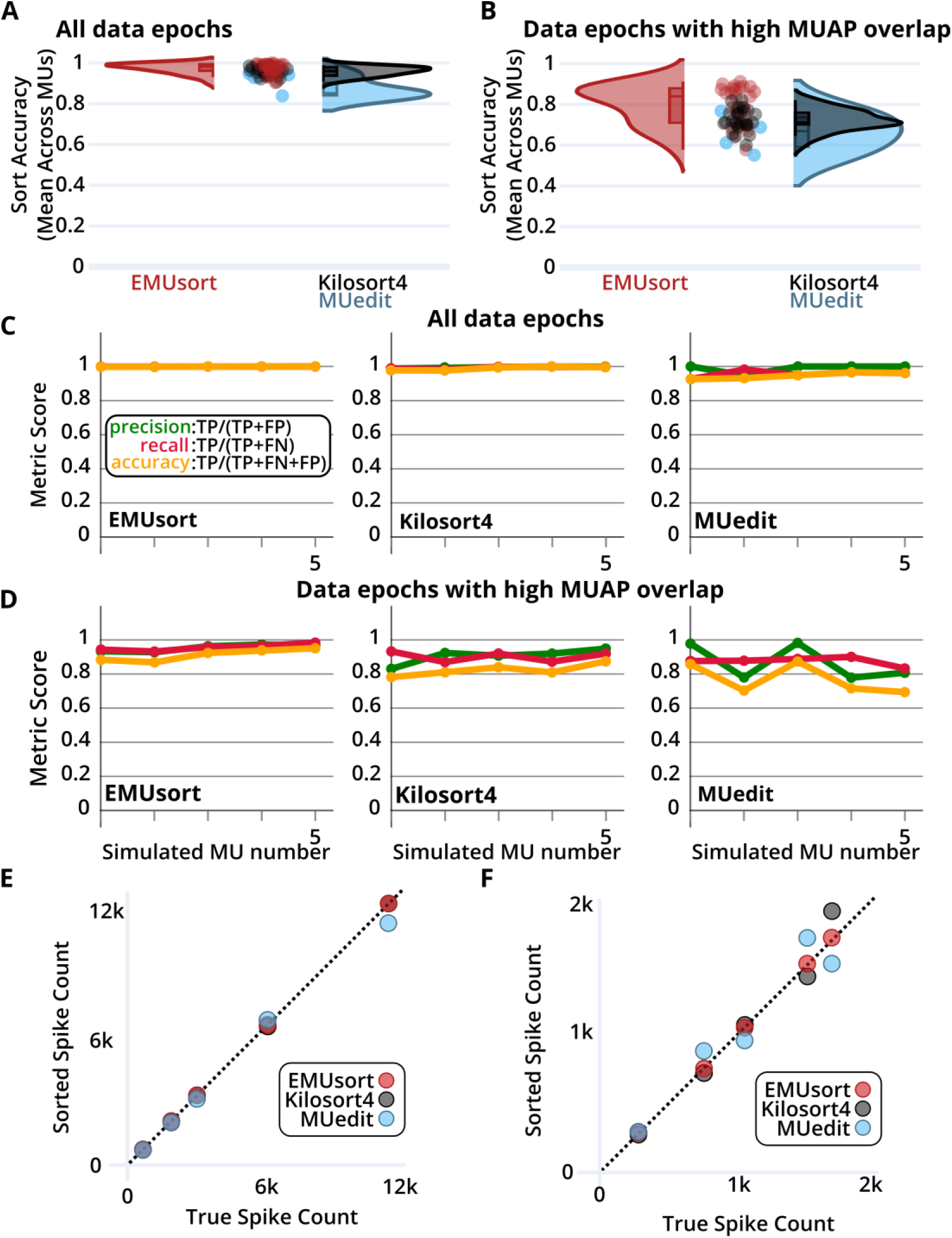
EMUsort achieves high performance with monkey-based simulations using real MUAP recordings from biceps brachii during reaching. (a) Graph of sort accuracy distributions, for EMUsort and KS4 across a parameter sweep with 25 combinations. Sort accuracies from four MUedit runs are also shown. Each point shows the mean accuracy across ground truth units for each sort run. (b) A similar graph as panel (a), but the accuracy calculations were computed only for highly overlapped spikes, i.e., only MUAPs within at least 2 ms of another MUAP were included in the calculations. This assesses how each sorter performs in the most challenging examples in the simulated dataset. In (a, b), the distribution plot is created with kernel density estimation for visualization. (c) Per-unit performance metrics from the best sort for each sorter using all spikes in the rat dataset, arranged from highest to lowest spike count. (d) Per-unit metrics for the same best sorts as in panel (c) but focuses only on overlapping spikes, as in panel (b). (e) Scatter plot showing how well the best sort from each sorter matches the true spike counts, where paired counts for each ground truth unit are shown as a point. The dotted line indicates where the sorted count should match the true count. Please note that panels (e, f) only reflect spike counts, and do not reflect accuracy. (f) A similar scatter plot, where MUAPs are only counted if they overlap with other MUAPs.

EMUsort showed a greater increase in variance for the overlapping results compared to the difference observed for the rat simulation (see Figure 6B, Figure 7B, and Tables 1–2). However, depending on the parameters used, a given EMUsort spike sorting run either outperformed or performed comparably to the other sorters. Compared to the rat simulation, this result could be driven by the lower total spike counts or the larger waveform width (4 ms versus 2 ms), which can both make proper template modeling more challenging. The higher variance of EMUsort in this case was skewed toward higher accuracies, producing the top 14 mean accuracy scores (out of 54 total mean accuracy scores) across all sorters with a single parameter sweep (see Table 4 for parameters used).

As in our analysis of simulated rat EMG data, we also compared the individual best sort run scores for simulated monkey data. EMUsort again produced the highest accuracy sort run for all simulated motor units with all spikes included (Figure 7C) and for overlapping spikes (i.e., MUAPs within half the spike width, 2 ms, of a neighboring MUAP; see Figure 7D). One difference from the rat simulation analysis (Figure 6) was that no motor units were completely missed in the best sort run for any of the sorters. With fewer simulated motor units and sparser overall activity (fewer total MUAPs; see Figure 7E-F), detection of all unique motor units was evidently easier to achieve for the monkey simulation. Driven by the higher accuracy scores, we see EMUsort achieved the closest match with the true spike counts both with all spikes included in (Figure 7E) and with overlapping spikes in (Figure 7F). KS4 and MUedit both exhibited higher deviations from the diagonal, showing higher or lower spike count values compared to the true simulated spike counts. Detailed per-unit statistics are presented in (Table 2 - table supplements 3-8).

For both datasets, an alternative accuracy score (as used in (Pachitariu et al., 2016)) and corresponding error rates were computed for all sorters (shown in Table 1 - table supplements 1-2 and Table 2 - table supplements 1-2). The data in these tables show that EMUsort provided the lowest median error rates across sorters. To calculate the percent reduction in median error rates when using EMUsort, we divided the median error rate for EMUsort by the same value for the sorter with the next lowest error rate (KS4; Table 1 - table supplement 2). This division yielded the percentage of errors present in the EMUsort sorts relative to those in the KS4 sorts (using median error rates across 25 sorts; 15.9 / 48.9 ≈ 0.3252). This result was then subtracted into one (1 − 0.3252 = 0.6748), which showed the percent reduction in median error rate when using EMUsort with the rat dataset was 67.5%. To compute the error rate reduction when using EMUsort with the monkey dataset, the same procedure was followed using the values in (Table 2 - table supplement 2). The ratio of median error rates was again computed between EMUsort and the next lowest error rate sorter (KS4; 17.0 / 33.9 ≈ 0.5015). This value was then subtracted into one (1 − 0.5015 = 0.4985), which showed that the percent reduction in median error rate when using EMUsort with the monkey dataset was 49.9%.

## Discussion

High-resolution recording of motor units provides insights into spiking codes for complex motor behavior that are not available from conventional bulk EMG recordings (Sober et al., 2018). Single-unit EMG recordings have revealed the diversity of coding properties within a single muscle (Thomas et al., 2024) and the role of millisecond-precise spike patterns in shaping behavior (Srivastava et al., 2017). Moreover, combining spiking measures of central neural activity with single-unit EMG allows direct comparison of spiking codes across forebrain, cerebellar, and spinal networks (Kirk et al., 2024). Myomatrix arrays (Chung et al., 2023) and other high density intramuscular electrode arrays (Metallo et al., 2011; Muceli et al., 2015) provide unprecedented access to high-resolution muscle data across species and muscle groups, necessitating efficient sorting algorithms, much as the development of Neuropixels probes necessitated the development of the Kilosort pipeline (Pachitariu et al., 2016, 2023, 2024; Steinmetz et al., 2021). To create EMUsort, we therefore adapted KS4 to sort high-resolution motor unit datasets.

EMUsort’s success in sorting motor unit data rests on several key modifications made to KS4 to account for the very different statistics of EMG versus neural data. First, EMUsort accounts for temporal delays between spike arrival times at different electrode contacts (Figure 2), which are much larger than those observed in neural data. Second, by providing a larger number of temporal components for waveform templates (Figure 3), EMUsort accounts for the greater waveform complexity of motor unit action potentials (Figure 1C) compared to action potentials in the brain (Figure 1F). Third, improvements in the selection and refinement of initial (“simple”) templates ensures the detection of later-recruited motor units, a common failure mode when the unmodified version of KS4 is applied to EMG data. Examination of both empirical and simulated data demonstrates the utility of these methods by quantifying the improved performance of EMUsort compared to both KS4 and MUedit (Figures 5–7).

EMUsort opens several avenues for new studies that rely on robust spike sorting of motor unit data. These analysis tools we present could accelerate studies that use high-resolution EMG signals recorded Myomatrix or other thin-film EMG electrodes (Chung et al., 2023; Metallo et al., 2011; Muceli et al., 2022; Oldroyd et al., 2025) in studies of clinical conditions that disrupt motor function (Grison et al., 2025; Ting et al., 2021). Other lines of investigation might combine high-resolution motor unit recording with simultaneously-recorded neural data to examine the neural coding of motor commands (Amematsro et al., 2025; Kirk et al., 2024; London & Miller, 2013; Marshall et al., 2022) or examine the effects of targeted electrical or optical perturbations on motor activity (Miri et al., 2017). EMUsort’s performance advantage over existing methods (Tables 1–2 and supplements) can therefore accelerate current and future studies that use high-resolution EMG data to explore neural function.

## Methods

### Surgical implantation of Myomatrix arrays in rat triceps brachii

All surgical procedures were approved by the Institutional Animal Care and Use Committee at Emory University (IACUC protocol number: 201700525). We implanted a single Myomatrix array (RF-4x8-BVS-8, previously RF400) consisting of 32 electrode contacts across four array “threads”) into the triceps brachii muscle of Long-Evans rat using methods described previously (Chung et al., 2023). One forelimb was targeted per animal. Rats were anesthetized with isoflurane in an induction box and moved to the surgical table where a steady level of anesthesia was maintained with isoflurane at 2-3%. The hair covering the head and upper forelimb was removed and small incisions were made on the scalp and near the targeted triceps muscle. The four threads of the Myomatrix array were then routed subcutaneously to the forelimb and inserted intramuscularly one by one into the targeted structure. The lateral head of the triceps brachii was used in this study, but sometimes the long head and extensor carpi radialis were used for testing. The forelimb incision site was closed with 6-0 suture (AROSurgical Black Polyamide Monofilament) and 4 to 6 FST bone screws (#19010-01) were implanted into the skull for mechanical support of the connector by pre-drilling with 0.9 mm burr FST drill bits (#19009-09). Dental cement was spread on the skull, surrounding the base of the screws and bonding the Myomatrix array’s miniaturized connector (Omnetics Inc.) to the skull. The metal contact of the ground thread of the Myomatrix array was then attached to an exposed bone screw with a silver conductive paint. Another final layer of dental cement encased the entire assembly.

### Data collection and rat behavior during treadmill locomotion

The treadmill used in this study was designed for rats and had variable inclines and speeds (Columbus Instruments Modular Rat Lane and Modular Rat Controller). Beginning the day after a surgical implantation, we would collect recordings in 1-2 minute segments across a range of walking and trotting speeds (5-50 cm/s, based on literature values for walking and trotting (Cohen & Gans, 1975; Pereira et al., 2006; Richards et al., 2019; Wisløff et al., 2001)) or inclines (0-20 degrees, in increments of 5 degrees). We replicated the same sequence of speeds or inclines for a maximum of 60 minutes during the session. Throughout the training phase before surgery, animals were always acclimated to the different speed and incline conditions, paying careful attention to signs of stress.

During treadmill sessions, EMG signals were recorded using an Open Ephys Acquisition Board along with Intan RHD 16-Channel Bipolar-Input Recording Headstages (RHD2216). Animals were briefly anesthetized with isoflurane at the beginning to be able to more easily plug in the tethered wire from the acquisition board to the Omnetics connector on the head of the rat. After doing so, at least 15 minutes were allotted for the rat to recover and acclimate to the treadmill before starting any EMG recording. Walking behavior was recorded on 4 FLIR BlackFly S video cameras (BFS-U3-16S2M-CS) at 125 FPS, for precise motion tracking of the 4 limbs, nose and base of the tail during locomotion. This motion information was used for determining the step cycles, and we used the implanted forelimb kinematics as the input signal to drive the simulated population of motor units during creation of our simulated datasets.

### Recording from Rhesus macaque forelimb muscle and reaching behavior

All procedures described below were approved by the Institutional Animal Care and Use Committee at Western University (IACUC protocol #2022-0208). One male rhesus monkey (Monkey M, Macaca mulatta, 10 kg, 15 years old) was trained to perform a range of reaching tasks while seated in a robotic exoskeleton (NHP KINARM, Kingston, ON). As described previously (Scott, 1999), this robotic device allows movements of the shoulder and elbow joints in the horizontal plane and can independently apply torque at both joints. Visual cues and hand feedback were projected from an LCD monitor onto a semi-silvered mirror in the horizontal plane of the task and direct vision of the arm was blocked with a physical barrier. An injectable Myomatrix array (NP-1x32-XZI-10, previously NG100) was inserted percutaneously into the biceps brachii muscle as described previously (Chung et al., 2023), recorded at 30 kHz as 16 bipolar channels using an Intan RHD2216 headstage and Open Ephys software. Then, using his right arm, Monkey M performed a reaching task to a continuous series of targets placed randomly within a 10 cm by 10 cm grid centered on the shoulder and elbow angles of 32° and 72°, respectively. Each trial began with the presentation of a new goal target (0.8 cm radius) and simultaneously a juice reward for acquiring the previous target. The subject then moved to the next target immediately. On one third of trials, an unpredictable mechanical perturbation was applied at a random point between the two targets with a fixed magnitude and duration (0.25 Nm pulse, 100ms), but a uniformly random Cartesian direction. After entering the goal target, the subject had to remain in the target for a variable time (300-500 ms) to complete the trial. On alternating blocks of 200 trials, there was either one goal visible (+0 trials) or two goal targets visible (+1 trials), and the subject had to reach the targets in order of appearance. Targets were selected to not be visually overlapping, and all targets were visually identical (white circle). Hand feedback was provided at the fingertip as a cyan circle (0.3 cm radius).

### Generation of simulated MUAP datasets

In order to quantify the MUAP spike sorting performance, we developed a simulation library, MUsim (O’Connell, 2025), which enables creation of simulated spike times and multichannel datasets using learned templates from previously spike sorted biological data. For clarity, we first outline the simulation process here at a high level, then we provide further detail in the subsections below. For each simulation, we began with a biological dataset and performed several spike sorting runs (with either EMUsort or KS4) until we found a high-quality sort output with minimal observed errors and a relatively high number of MUAP clusters. This was a supervised process that relied on manual evaluation in the Phy GUI. We eliminated poor clusters based on presence of one or more types of error (type I, type II, impossibly high or dramatic changes in firing rates, multimodal amplitude distributions, etc.). Once we selected the clusters to include in the simulation, we used the corresponding spatiotemporal templates derived from Kilosort that represented each putative motor unit. These templates have temporal components that define the range of shapes the spikes exhibit and spatial components, or channel weights, that define how much of each temporal component to apply at each channel. Temporal components, *W*, are defined by applying PCA to a set of non-overlapping waveforms as described in “Kilosort represents spike features using principal components from the non-overlapping spikes”. Channel weights, *U*, are defined for each cluster by convolving each temporal component at the corresponding spike times to produce weights of match strength for each temporal component in the form of PC loadings for each channel. In this way, the channel weights represent the waveform shapes on each channel for each putative motor unit, as described in “Kilosort classifies spike features with advanced clustering methods and spatiotemporal constraints”. Together, these temporal components and channel weights were used as multichannel templates to produce each simulated dataset. To model the variance in waveform shapes that we observed empirically, for each putative motor unit, we used the original set of temporal components along with noise randomly added to the channel weights for each spike example, producing a variable waveform at each spike time.

#### Matching waveform shape variability

To simulate the variability in waveform shape empirically seen in our electrophysiological recordings, we used the following strategy. If *k*, *t*, *c*, and *p* indicate the number of spikes, timepoints in each spike, channels, and PCs of the non-overlapping waveforms (see “Kilosort represents spike features using principal components from the non-overlapping spikes”), respectively, and 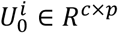 is a matrix of weights used to reconstruct the template for the *i*^*th*^unit from the PCs (also called temporal components), we create *U*^*i*^ ∈ *R*^*k*×*c*×*p*^by duplicating the same values of 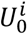 along the first dimension as placeholders for the number of spikes we will generate. Next, a matrix *G*^*i*^ ∈ *R*^*k*×*c*×*p*^; *G*^*i*^ ∼ *N*(*μ*, *σ*) is filled with samples from a Gaussian distribution to be the same shape as *U*^*i*^. Finally, we create *W* ∈ *R*^*p*×*t*^as a matrix containing the PCs. Using these matrices, we were able to generate unique MUAP spikes for the *i*^*th*^simulated unit and *k*^*th*^spike instance in the following way:

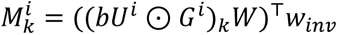

Where 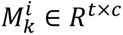 is the unique “morphed” MUAP waveform for the *i*^*th*^ unit and *k*^*th*^ spike instance, *b* is a positive scalar to convert from bits into voltage units, and *w*_*inv*_ ∈ *R*^*c*×*c*^is the inverse whitening matrix used to undo the channel whitening performed in KS4. The ⊙ operator denotes element-wise multiplication of two equally sized matrices. For *G*^*i*^, μ=1 and σ=0.2, with the standard deviation level chosen empirically to match the level of shape variability within each cluster to the observed variability in the real data.

To ensure the edges of the waveforms are brought to zero (i.e., to prevent discontinuities at the edges of each spike example), MUAPs were windowed using the following procedure: If *D* ∈ *R*^*t*×*c*^ is a data matrix to contain all multichannel MUAPs *M* applied additively at each spike time *T*^*i*^ ∈ *R*^*t*^, the MUAPs in *M* of width *n*_*t*_ (2 ms and 4 ms for the rat and monkey datasets, respectively) are windowed in *D* by multiplying a zero-padded Tukey window *t*_*w*_ with *α* = 0.25, window width 0.9*n*_*t*_, and zero-padding of width 0.05*n*_*t*_ before and after (total width *n*_*t*_).

This approach yields a multichannel spike train for each simulated unit, with variability in the shape for each spike instance, consistent with our biological recordings, but lacking any of the observed baseline Gaussian noise.

#### Matching channel noise levels

To ensure that each channel of the simulated time series had similar noise levels to the original data *D*_0_ ∈ *R*^*t*×*c*^, we first computed the standardized median absolute deviation (MAD) values for each channel of the original data:

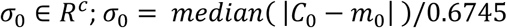

Where *C*_0_ ∈ *R*^*t*×*c*^ is a chunked subset of *D*_0_, *m*_0_ ∈ *R*^*c*^ are the medians of each channel of *D*_0_, and 0.6745 is a scaling factor to produce an estimate of the standard deviation, for Gaussian distributed data (Magland et al., 2020; *SpikeForest*, 2025).

The following optimization procedure was then used to fit the noise level of each channel to the original recording, after the MUAPs *M* were added into *D* at times *T* and windowed by *t*_*w*_. Standard deviation values for additive Gaussian noise on each channel are updated through gradient descent using the Adam optimizer from Pytorch (Kingma & Ba, 2017; Paszke et al., 2019). An array of standard deviations *σ*_*L*_ ∈ *R*^*c*^ is the only learnable parameter, where learning continues until MAD values for all channels are within 1% of the original noise level for each corresponding channel. For each iteration, a new *D*_*aug*_ ∈ *R*^*t*×*c*^is computed with the updated *σ*_*L*_ array and *σ*_*S*_ ∈ *R*^*c*^is defined as the MAD of *D*_*aug*_ as below:

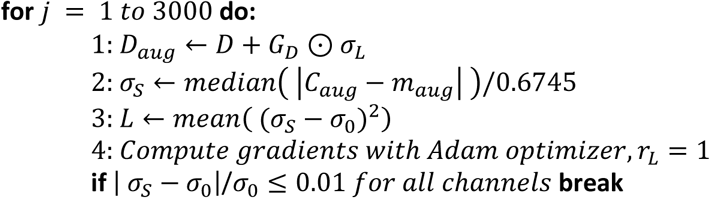

Where *D*_*aug*_ is the simulated data *D* after adding Gaussian noise to each channel, *σ*_*L*_ is a learned array of different standard deviation values to independently fit each channel noise level, and *G*_*D*_ ∈ *R*^*t*×*c*^; *G*_*D*_ ∼ *N*(*μ*, *σ*) with μ=0 and σ=1, *C*_*aug*_ ∈ *R*^*t*×*c*^ is a chunked subset of *D*_*aug*_, *m*_*aug*_ ∈ *R*^*c*^ are the medians of each channel of *D*_*aug*_, *L* is the mean squared error loss function, and *r*_*L*_ is the learning rate. The ⊙ operator denotes element-wise multiplication along the broadcastable dimension.

#### Spike times driven by recorded kinematics with MUsim

Using MUsim, the spike times were generated for each motor unit based on previous models of motor unit recruitment and firing dynamics (Fuglevand et al., 1993b; Heckman & Enoka, 2012; Hennig & Lømo, 1985). An object, *M*_*sim*_, contains all simulation parameters, “sampled” unit information (e.g., activation thresholds), applied forces, and the corresponding output spike trains for each simulated motor unit. If the number of motor units, *u*, is set by the user, and if 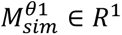 are scalars acting as lower and upper bounding parameters for an exponential distribution, the activation thresholds of each unit are defined as 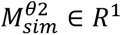 where *λ* = 1. Once these thresholds are sampled for each unit, a force (or kinematic) array can be passed into the object as 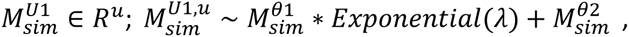 where *t* is the time dimension. This operation results in a different response curve from each unit, 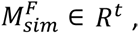 which is initialized for each unit as the subtraction of the activation threshold into the force signal:

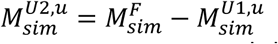

These response curves control the probability of a spike occurring for each unit and each time sample, and they are continuously updated whenever a spike occurs by adding a spike history time kernel, 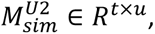 to greatly reduce the probability of another spike occurring immediately after a spike event (Pillow et al., 2008). This spike history kernel precisely controls the max firing rate with precise spike timing and prevents refractory period violations. For each time point, the spike events, 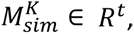 are determined by passing the response curves through a sigmoid and drawing a sample from a binomial distribution according to the following algorithm:

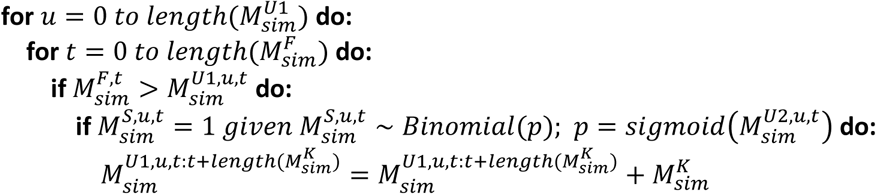

In practice, the units are computed in parallel, whereas the time loop must be computed in series because all loop iterations depend on the result from the previous iteration. When this algorithm completes execution, all spike times are defined for all units. In our simulations, the firing rates are coordinated in concert with fluctuations of the kinematic signals we recorded during treadmill locomotion using DeepLabCut and Anipose with 4 camera triangulation (Karashchuk et al., 2021; Mathis et al., 2018), sampled at 125 FPS.

In this way, we created a 10 minute simulated dataset with 10 different experimentally realistic waveform shapes across 8 channels, which matched the number of good Myomatrix electrodes implanted in a rat triceps brachii during unrestrained treadmill locomotion. We also used this method to create a 10 minute simulated dataset from 5 experimentally realistic waveforms taken from Myomatrix recordings in the biceps brachii of a monkey during a random target reaching task. We then used these known spike times for each simulated motor unit as the ground truth to assess spike sorting quality for the spike sorters being compared, which is discussed below.

### Evaluating sort quality with simulated MUAP datasets

For performance evaluation, we follow a two-step process: we first match putative units identified by a sorter to the closest corresponding ground truth unit. Next, for each pair of sorted and ground truth units we perform time alignment and determine the overall number of spike time matches in each matched pair, leading to objective scores for precision, recall, and accuracy corresponding to each ground truth unit. To do this, MUsim uses an efficient three stage procedure with increasing time resolution:

The first stage is greedy cluster matching. Its primary purpose is efficient pairing of the best matched sort clusters to each ground truth unit, so it is conducted with coarse temporal resolution (10 ms bins). Spike time data are first binned into 10ms bins, so the spike times are represented by the number of spikes for each cluster in each bin, in a large *N* x *T* matrix (where *N* is the number of clusters and *T* is the number of time bins). This matrix is then used to match clusters using the accuracy metric as the pairing criteria, where the best clusters are paired with ground truth units without replacement in descending order of accuracy.

The second stage performs alignment at high temporal resolution using binned spike arrays (0.1 ms bins) for each matched cluster based on correlation maximization with the ground truth binned spikes. This procedure is necessary because a sorter can define spike times for any phase of a MUAP, whereas the simulated ground truth times are defined at the center of each template. The correlations are computed for each matched pair, between the ground truth binned spikes and time-shifted copies of the binned spikes for each matched cluster within +/- 2 ms (41 copies between +/- 20 bins). Whichever time shift produces the highest correlation for each matched pair is used for the final spike time matching stage.

The third stage uses the same high time resolution (0.1 ms), this time operating on spike times (instead of binned spike arrays in the previous stage) to check for spike time matches with the best shifted array for each matched pair. To do this, the 0.1 ms bins are converted into spike times by finding the indexes of all bins containing a spike. A vectorized algorithm then subtracts each ground truth spike time into the sorter spike times array followed by taking the absolute value, which allows thresholding by 1 ms to quantify the number of sorter spike times that match within +/- 1 ms. The algorithm is careful to count the number of refractory period violations, where multiple spikes could occur within the +/- 1 ms window. In this case, only the closest spike is taken as the true positive, and the remaining spikes are counted as false positives, because we enforced that refractory period violations could not occur during creation of our ground truth arrays.

We next compute standard classification performance metrics (as shown in the equations below) based on the spike time matches. The first set of equations defines true positives, false positives, and false negatives, and the second set shows how three standard performance metrics (precision, recall, and accuracy) are computed (Magland et al., 2020).

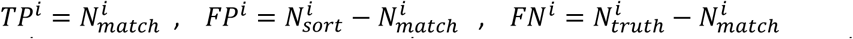

Where *TP*^*i*^ is the count of true positives, *FP*^*i*^ is the count of false positives, and *FN*^*i*^ is the count of false negatives for the *i*^*th*^ matched pair.

These equations show the equivalence with 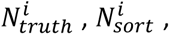 and 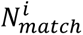 that we use in our calculations, which are counts of ground truth spikes, sorter labeled spikes, and spike time matches, respectively, for the *i*^*th*^ matched pair.

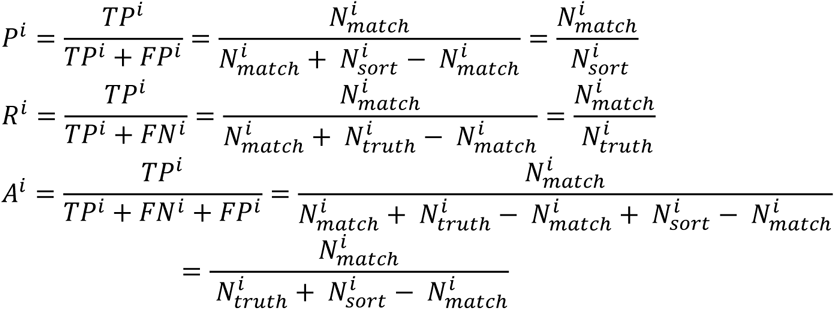

Where *P*^*i*^, *R*^*i*^, and *A*^*i*^ are the computed precision, recall, and accuracy for the *i*^*th*^ matched pair. See (Supplementary Figure 3) for examples of each error type designation when spike sorting simulated spike trains with EMUsort and KS4.

### Considerations in the approach for performance evaluation

There are several decisions to make prior to being able to compute the performance of a spike sorting run that greatly influence the resulting score. The current approach conservatively quantifies the performance of sorting using the accuracy metric defined in “Evaluating Sort Quality with Simulated MUAP Datasets” in Methods, which accounts for both false positives and false negatives. Before this accuracy calculation, we had to make a supervised decision to determine the number of ground truth motor units to evaluate against, although this is sometimes ambiguous, depending on the dataset. During generation of the simulated datasets, we determined the number of ground truth units to simulate by manually evaluating a sort run using several quantitative methods such as cross-correlogram charts to assess refractory period violations, amplitude distribution Gaussianity to assess contamination (false positive content), and inspecting the time series data at random intervals across the dataset to check for obvious classification errors. This was how we determined the number of ground truth clusters to evaluate against. In the future, this approach could be automated through use of composite scoring to remove the subjective nature of this evaluation. In many datasets, there can exist several MUAPs that are too small to be classified robustly due to low SNR or low amplitude relative to the other larger MUAPs that tend to coactivate, causing overlaps that inject variability larger than the size of the smaller MUAP so that it cannot be well isolated, even after using deconvolution approaches. In cases like these, if these motor units had been simulated, the accuracy for those ground truth motor units would be close to zero, which would greatly reduce the overall median accuracy score.

### Parameter sweep comparisons

Using the approach described in “Generation of simulated MUAP datasets” to create simulated datasets, we ran multiple spike sorting runs for both KS4 and EMUsort across a range of parameters related to 1) threshold crossing levels (in units of standard deviations) and 2) similarity thresholds for determining spike cluster assignments. The parameter sweeps across KS4 and EMUsort were kept identical except in the case of the single channel threshold for getting threshold crossings (*Th_single_ch*) because EMUsort enables aggregation of spikes across multiple thresholds whereas KS4 only allows a single threshold crossing at a time. To minimize this difference during the parameter sweep, we made the single thresholds for KS4 span the same range as all the thresholds included for EMUsort. We also tried several different threshold sweeps for KS4 and proceeded with the set that produced the best results for KS4 for the final analysis. For the parameter sweep results shown in this paper for KS4 and EMUsort, we swept the parameters shown in (Tables 3–4) to produce 25 different sorts for each.

Because of the differences in the implementation of MUedit, we could not perform parameter sweeps in a directly comparable way. Instead, we first set base parameters that were in line with recommendations given by the authors, such as enabling Peeloff and disabling the CoV filter. For the rat simulated dataset with 8 channels and 10 motor units, we set the number of extended channels to 80 and for the monkey simulated dataset we chose 160. The number of iterations for all runs was set to 160 and we always used 1 time window that included the full 10 minute dataset. We explored 4 combinations of the initialization and contrast function parameters, which were roughly comparable to the *Th_single_ch* setting (which affects initialization) and together as linked parameters, *Th_universal* and *Th_learned* (which both affect matching).

### Producing composite scores for agnostic estimation of sort quality

With EMUsort, it is possible to run a large number of processes in parallel, which was intended to save the user time, but considering the standard manual process of cluster evaluation, it would be even more labor intensive to check across all sort outputs and evaluate which is best. So, we needed to provide a method of automatic evaluation of an entire sort run, so that a user could run a wide parameter sweep followed by a manual evaluation of only the best performing sort runs. To do this, our goal was to emulate the standard manual process of evaluating each cluster by using several metrics of error ranging from 0 to 1 and multiplying them together to aggregate the results into a single value so that the presence of a single type of error in a cluster would result in a bad composite score. Afterwards, these composite scores for each cluster could be averaged to provide an overall assessment of sort quality for each sort run.

To build a composite metric like this, we took advantage of existing metrics that are publicly available in the quality metrics module from SpikeInterface to quantify these different types of error. The functions will be explained further below, but for clarity, we used their implementations to compute the following metrics for each cluster: the Llobet contamination ratio, presence ratio, amplitude cutoff, firing rate, firing rate range, and SNR. We augmented and combined some of these metrics to allow them to meaningfully range from 0 to 1 while representing a given error type. Some of the terms already ranged from 0 to 1, so all we had to do was reverse the polarity by subtracting into 1. All augmentations are indicated in the below equations, but the metrics themselves are defined elsewhere (Buccino et al., 2020).

The composite score we use has 4 components for 4 different types of error: Type I error (false positives), type II error (false negatives), invalid firing rates, and low SNRs. This design was chosen so that the scores for each error type could be multiplied together to emulate the process of elimination where the presence of a single error results in rejection of a cluster. This structure results in aggressive penalties for a cluster if multiple types of error are present at the same time, forming a highly conservative estimate of overall cluster quality. After composite scores are computed for each cluster identified by the sorter, they are averaged together to represent the overall proportion of good clusters in a single conservative value, ranging from 0 to 1. A high overall composite score for a sort is therefore only possible if there is a low number of bad clusters, leaving mostly clusters that have high composite scores. Below the construction of each score component is detailed.

The type I error component of the score estimates the fraction of false positives in a spike train. This is calculated with the SpikeInterface implementation of the Llobet contamination ratio to produce a measure of contamination, which is then subtracted into 1 so that a score of 1 means no type I error detected.

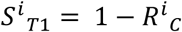

Where *S*^*i*^_*T*1_ is the type I score for the *i*^*th*^ cluster and *R*^*i*^_*C*_ is the Llobet contamination ratio for the *i*^*th*^ cluster with a 1 ms refractory period (Buccino et al., 2020; Llobet et al., 2022).

The type II error component captures information about false negatives using the SpikeInterface functions for computing presence ratio (20 second bins, and 0.5 mean firing rate threshold) and for amplitude cutoff (32 bins) which estimates the fraction of how many spikes are missing based on an assumption of Gaussianity of the amplitude distribution for a valid MUAP cluster (Buccino et al., 2020).

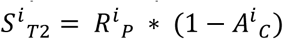

Where *S*^*i*^_*T*2_ is the type II score for the *i*^*th*^ cluster, *R*^*i*^_*P*_ is the presence ratio for the *i*^*th*^ cluster, and *A*^*i*^_*C*_ is the amplitude cutoff for the *i*^*th*^ cluster.

The firing rate validity score component simply assesses whether a given cluster exhibits plausible firing rates for a motor unit, across common model animals, and essentially removes a given cluster from consideration (brings composite cluster score quickly toward 0) if the mean firing rate rises above 200 Hz. It also penalizes unreasonably large changes in firing rate (>200 Hz) by incorporating the SpikeInterface function for firing rate range, which computes the difference between the 95th and 5th percentile firing rates with a bin size of 0.5 seconds.

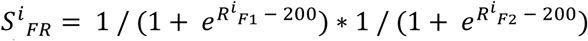

Where *S*^*i*^_*FR*_ is the firing rate validity score for the *i*^*th*^ cluster, *R*^*i*^_*F*1_ is the firing rate for the *i*^*th*^ cluster, and *R*^*i*^_*F*2_ is the firing rate range for the *i*^*th*^ cluster.

The SNR score component is another sigmoidal score that heavily penalizes a cluster if it goes below 4 SNR. The decision of 4 SNR as the midpoint (produces a value of 0.5) was based on our own observations that sorting is generally better above this level. This observation is strengthened by the result shown in Figure 3 of the 2020 SpikeForest paper, which shows 4 SNR to be the inflection point for higher ground truth accuracies in a Kilosort2 analysis (Magland et al., 2020).

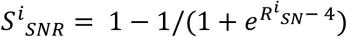

Where *S*^*i*^_*SNR*_ is the SNR score for the *i*^*th*^ cluster, and *R*^*i*^_*SN*_ is the SNR of the *i*^*th*^ cluster.

For each cluster, the EMUsort composite score is the product of the above 4 score components.

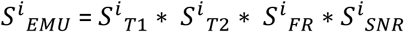

Where *S*^*i*^_*EMU*_ is the EMUsort composite score for the *i*^*th*^ cluster.

The overall EMUsort composite score for a given sort is the mean across cluster composite scores.

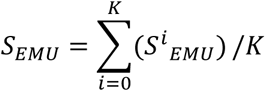

Where *i* is the cluster index and *K* is the total number of clusters identified for that sort run.

This way, if a sort were to have a high proportion of high-quality clusters with low contamination, high recall, reasonable firing rates, and SNRs above 4, that sort would receive a higher composite score relative to other sorts with a lower proportion of high-quality clusters. If a sort has good clusters but also a higher number of bad clusters, the mean score is brought down, which is desirable because a sort like this places a higher burden on the user to inspect and prune through the larger number of bad clusters. The mean across clusters was chosen over the median because the mean of multiplied scores across units was found to produce higher correlation with ground truth accuracy than median scores across units. By design, the overall structure of each component of the composite score was not considered critically important as long as each effectively achieved the goal of containing smoothly ranging information about the different error types. This composite scoring method can likely be further improved in future work, but to our knowledge, EMUsort is the first publicly accessible spike sorting tool to provide parameter sweeping with composite scoring to estimate the quality of entire sort runs, which is something highly valuable to a user who wants good sorting results as quickly as possible. Overall, EMUsort uses publicly available functions to create a novel composite score across all units, and the score correlates positively with ground truth accuracy for multiple datasets (Supplementary Figure 2).

### Main configuration file controls when using EMUsort

When EMUsort is executed in the terminal, the main, five-part configuration file can be adjusted, and the settings it contains are referenced throughout the entire pipeline. For a complete list of the parameters, see Table 5. First, the Data section plays a role in how the dataset is loaded and preprocessed: what data format to expect, which experiments to use, whether to concatenate multiple recordings, what bandwidth to use for bandpass filtering, and whether to slice the time of the recording.

**Table 5:**
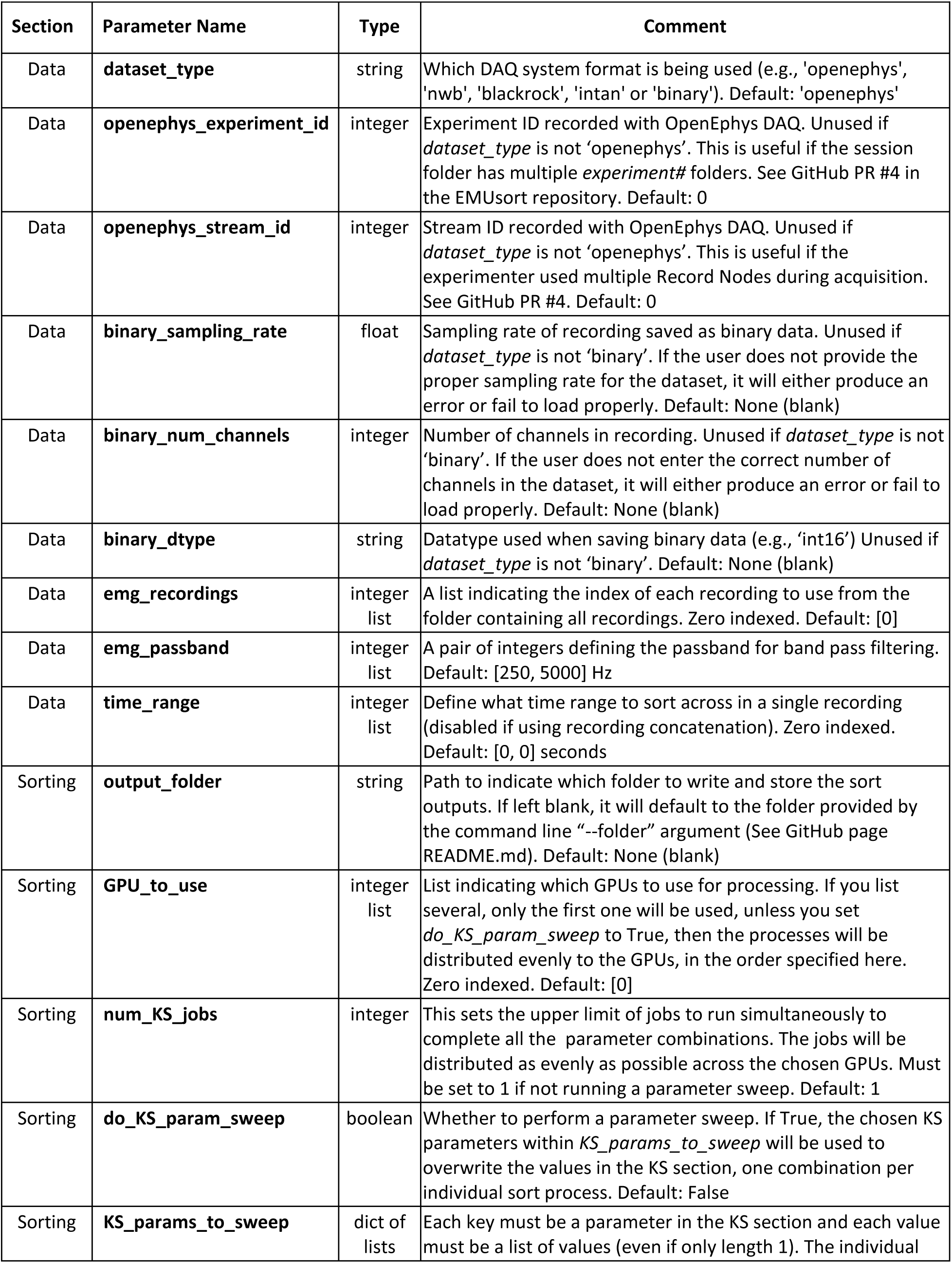
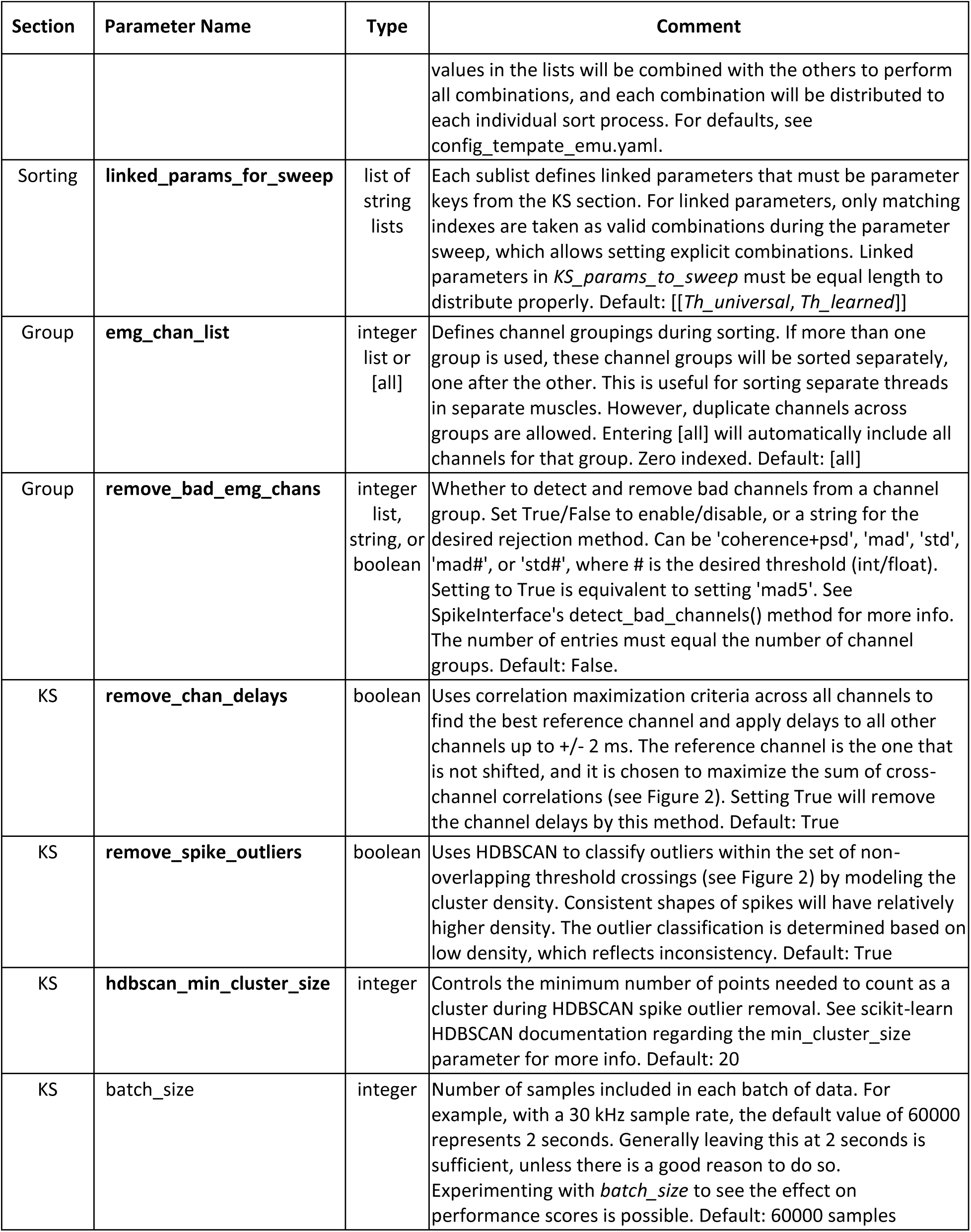
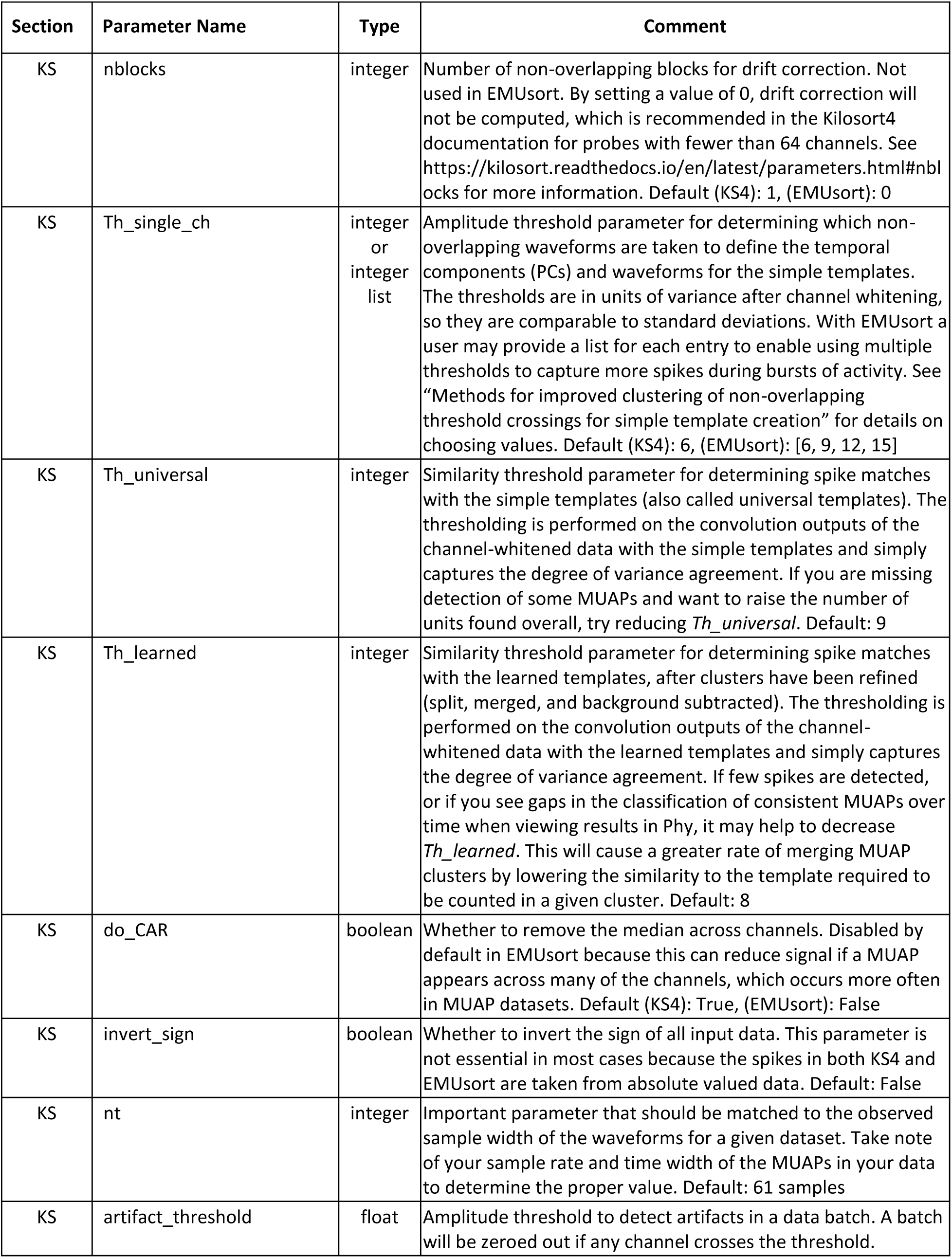
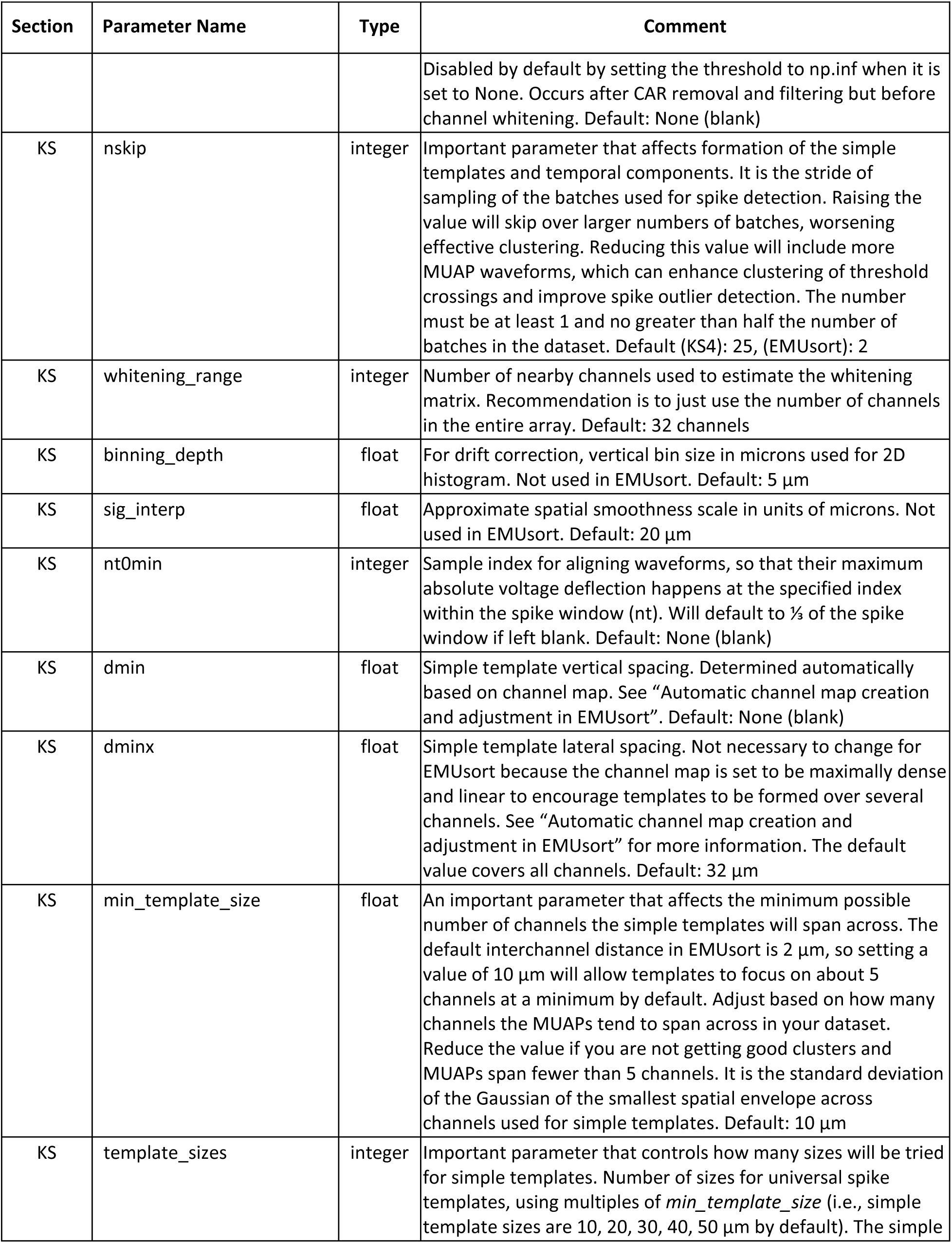
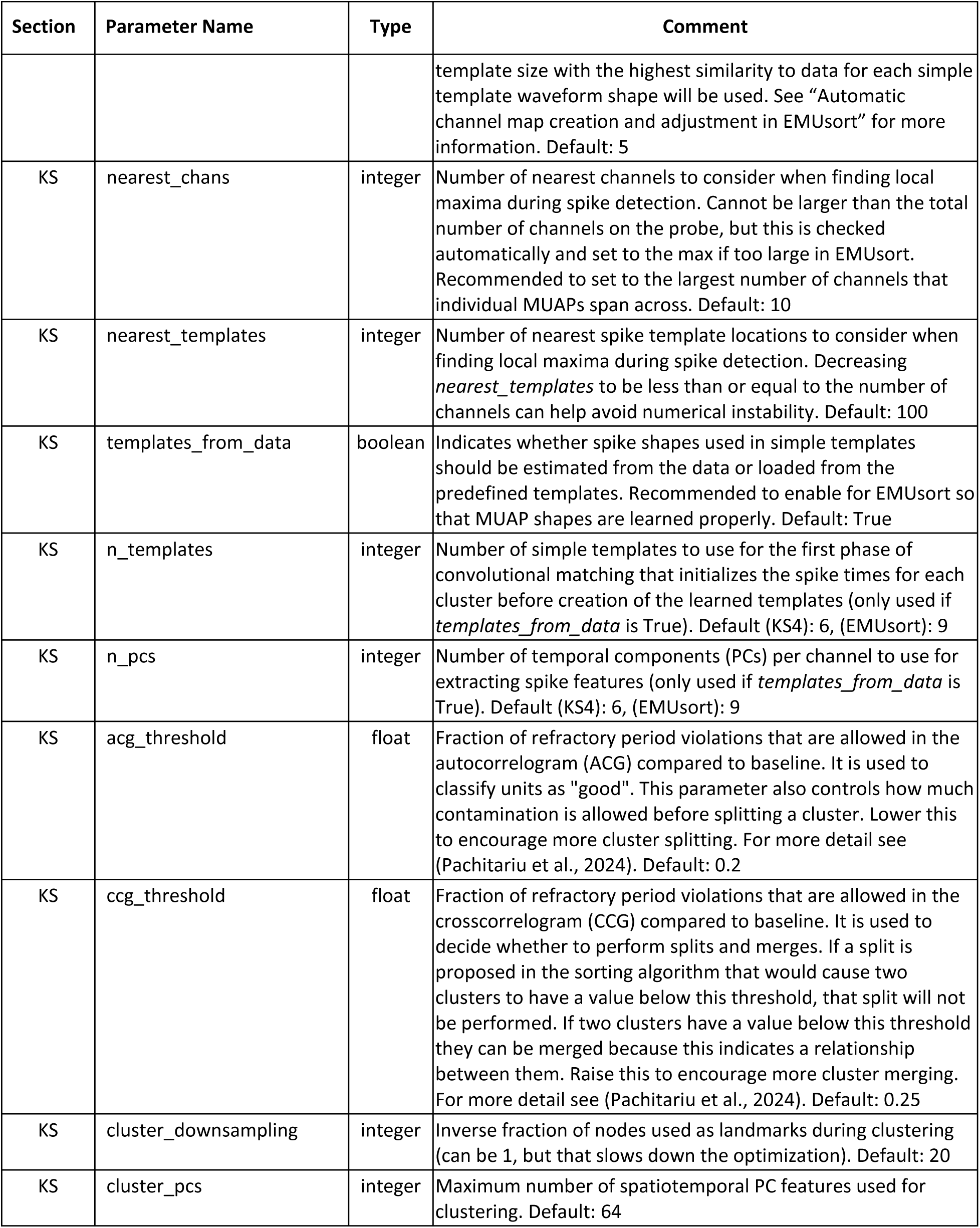
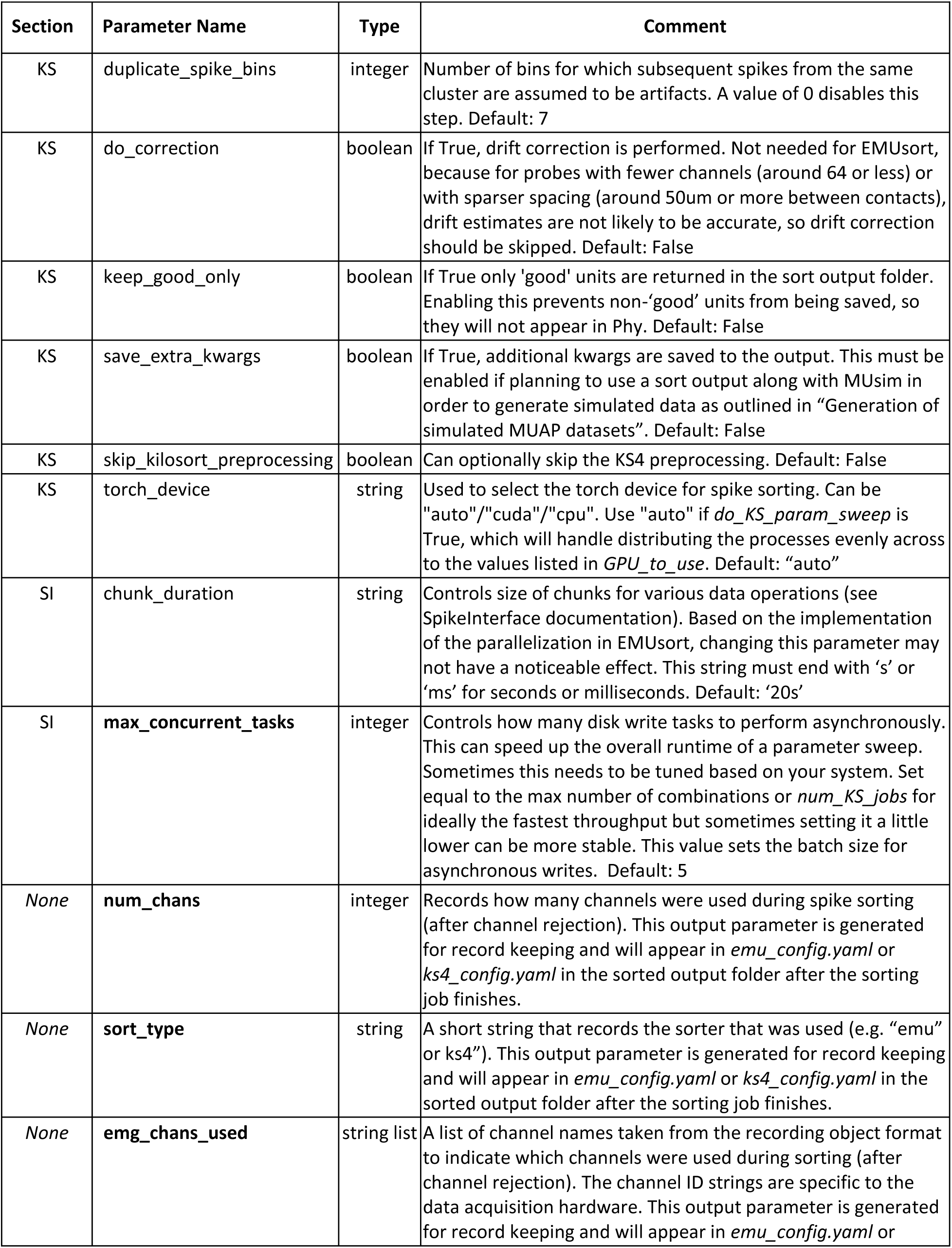
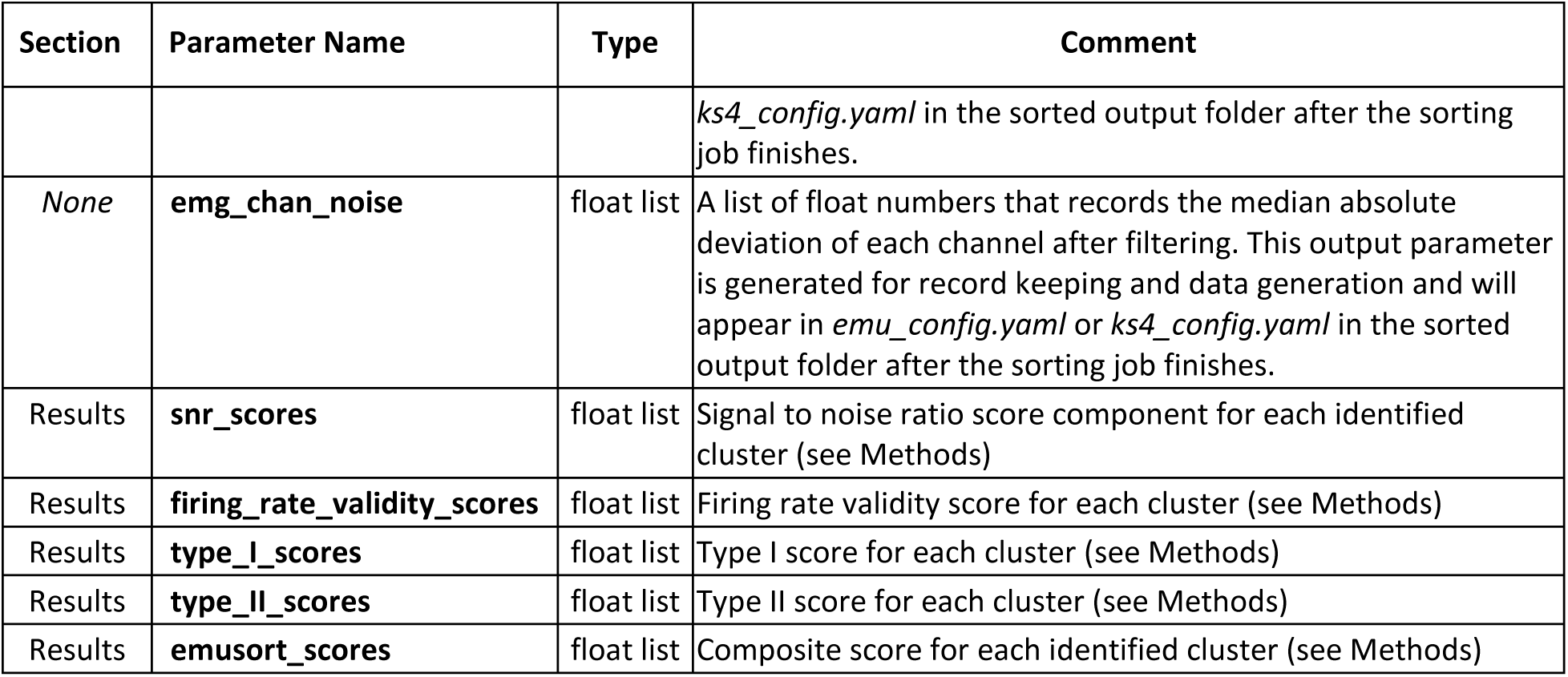
EMUsort configuration parameters. List and explanation of important EMUsort parameters that control its operations. Bolded parameters were introduced in EMUsort, whereas items that are not bolded already existed in KS4, but are made accessible in the EMUsort configuration file for greater usability and parameter sweeping. For shared parameters, default values are for both EMUsort and KS4, unless otherwise noted. Note that the parameters marked “None” or “Results” for the Section field only appear in the copy of *emu_config.yaml* automatically saved to a sort output folder after a sort run is completed and are simply appended to the bottom for record keeping and usage in future analyses.

Second, the Sorting section controls what folder the sort outputs will be sent, which graphics processing unit (GPU(s)) to use, whether to perform a parameter sweep, and the main threshold parameters of KS4.

Third, the Group section allows the user to manage which channels are sorted. If multiple groups of channels are specified, each channel group will be sorted separately. For each group, the user can set the method by which bad channels will be detected and removed. The Group section allows running multiple sorts sequentially with different combinations of channels as a group to be sorted together. For example, a user could set *emg_chan_list* as [0,1,2,3] for one thread implanted into one muscle and [4,5,6,7] for the second group in a different muscle. The number of Group parameters must match to be properly distributed. If two channel groups are specified, the user also needs to specify two entries for the bad channel removal parameter, *remove_bad_emg_chans*, such as [True; True]. A user could also set a single group as [0,2,3] if channel 1 is known to be broken, and [False] for *remove_bad_emg_chans* to disable automatic channel rejection. Setting multiple channel groups is compatible with the parameter sweep function.

Fourth, the KS section controls all KS4 parameters, which are documented elsewhere (https://kilosort.readthedocs.io/en/latest/parameters.html), but are also outlined in Table 5. This section contains three additional EMUsort-specific parameters that control 1) whether time delays should be removed from channels by the method described in Results (*remove_chan_delays*, see Table 5), 2) whether outlier spikes should be removed during initialization of the templates (*remove_spike_outliers*, see Table 5), and 3) the minimum number of spikes needed to be considered a cluster for HDBSCAN during the outlier removal stage (*hdbscan_min_cluster_size*, see Table 5).

There is a final minor section called SI which allows the user to control parameters related to SpikeInterface and how the sorting results are batch processed. The *max_concurrent_tasks* parameter (see Table 5) allows controlling how many sort output folders to write concurrently, which can greatly speed up saving. Sometimes there are system limitations on the total number of files that can be opened per process, so to avoid an error in Linux systems, EMUsort raises this resource limit if it is too low.

There is another configuration file (*ks4_config.yaml*) that is generated if the user performs an emulation of default KS4 using EMUsort commands (see documentation on the main Github page, https://github.com/snel-repo/EMUsort). This configuration file mirrors the same structure of *emu_config.yaml* but it sets all defaults as found in the main KS4 repository and disables all EMUsort procedures so that the user can easily test for performance differences, if desired. Because this feature is designed for KS4 emulation, the *ks4_config.yaml* file is reset for each run and does not interfere with the main configuration file.

Overall, the main configuration file (*emu_config.yaml*) controls different stages of the pipeline, and if the parameter sweep option is enabled in the Sorting section, EMUsort uses SpikeInterface methods to make parallel calls to the Kilosort4Sorter class. Separate configurations are then copied and sent to each subprocess with each unique parameter combination, and EMUsort distributes the jobs equally amongst the available GPUs for fastest completion. By using this method, several parameter combinations can be tried during spike sorting of a given recording in roughly the same amount of time it would take for a single run.

### Innovations in the development of EMUsort and usage tips

EMUsort builds upon existing open-source libraries to substantially improve spike sorting performance for high-resolution EMG datasets across a range of parameters (Figure 6A-B) while also providing a command line interface with high usability and a novel combination of functionality (batch runs with channel grouping, parameter sweeping parallelized across multiple GPUs, sorting performance estimation, etc.; see Supplementary Figure 1). EMUsort allows users to run sorts in parallel across a range of sorting parameters, which are then scored using the composite score described in “Producing Composite Scores for Agnostic Estimation of Sort Quality”, which produces an estimate of the most likely best performing sorts. When sorting biological datasets, the user cannot access ground truth to compute a true accuracy score, so EMUsort provides this composite score as a quantitative aggregate metric that serves as a proxy for accuracy.

When we evaluated the true accuracy scores for parameter sweeps with 2 different simulated rat datasets, we found the composite scores were positively correlated with the mean ground truth accuracy scores (Supplementary Figure 2). For both parameter sweeps, the most objectively accurate sorts were among the top 5 sort runs with the highest composite scores. Therefore, it appears to be good practice when using the EMUsort parameter sweep functionality to check the sorts in descending order of the EMUsort composite score. By following this procedure, a user can ideally save time by only evaluating the top sort runs (e.g. the top scoring 5 out of 25), while likely attaining a better final sorting result compared to running only 5 sort processes sequentially, without the sort quality estimation.

### Preprocessing stages in EMUsort

EMUsort performs a range of automatic and adjustable preprocessing steps that help to improve performance of downstream spike sorting operations. All parameters used for each run are dumped into each sort folder inside the *emu_config.yaml* file, which serves as a record of all user settings for each sort run.

Recording concatenation allows finding MUAPs across multiple recordings. Which recordings are concatenated is adjustable in the EMUsort configuration file. The user must keep track of recording lengths and the order that recordings were used in to be able to reconstruct the true spike times later during analysis against behavioral or task signals. The order of recordings can be found in the dumped *emu_config.yaml* file in the sort output folder.

Automatic noisy channel removal can be performed with Spikeinterface methods if enabled. The user can specify different threshold levels in the configuration file (with median absolute deviation or standard deviation). Lower values cause more channels to be rejected, which can improve sorting if those channels were too noisy, contained artifacts, or didn’t have significant shared activity with other good channels.

Bandpass and notch filtering is performed with standard preprocessing operations using SpikeInterface functions. The bandpass filter is adjustable in the configuration file and helps focus the downstream analyses on information in the spike band of 250-5000 Hz. The notch filter is set by default for the United States value of 60 Hz to remove line noise and would need to be adjusted in the code by the user to match the line noise level to benefit from it in other countries (e.g., 50 Hz).

The time delay removal across channels is also performed as a preprocessing step alongside the whitening operations within KS4. The function we implemented, *get_channel_delays*, mirrors the implementation of the *get_whitening_matrix* function in KS4 so that it finds the correlation-maximizing delays to apply for each channel at the beginning, stores those delay values for each channel as an array, and simply applies those shifts to each channel upon the loading of each batch. One difficulty with computing a general set of shifts to apply for alignment of all MUAPs is that the muscle fibers for each motor unit are arranged uniquely within the muscle, so each motor unit could theoretically have a different set of optimal channel shifts. By applying our method, we essentially take a linear fit between the best channel shift values for each motor unit, weighted by the integral of the signal contributions from each motor unit, so that motor units with larger amplitude or larger spike count would result in a larger influence on the final set of channel shifts that are computed by correlation maximization. Afterwards, each time a new batch of data is loaded (2 seconds by default), channels are shifted in time by the previously quantified amount to minimize the MUAP time delays (Figure 2B).

Finally, because all Myomatrix devices currently have fewer than 64 channels, we disabled drift correction in EMUsort by setting *nblocks* to 0 by default (see Table 5). This conforms with KS4 documentation to avoid inaccuracy in the drift estimation (Pachitariu, 2024).

### Automatic channel map creation and adjustment in EMUsort

EMUsort automatically generates a dense, linear channel map of minimal size (2 μm between channels) for convenience and to make KS4 include more channels in the creation of templates. We want this behavior because MUAPs appear across many channels due to the way the depolarization spreads across muscle tissue. To control how many of the channels are used in the templates, the *min_template_size* can be adjusted. We left the *min_template_size* parameter as the default value of 10, so that it will use channel weights to focus on a minimum of 5 channels during convolution of the simple templates. If there are some MUAPs that don’t appear on as many channels, it might be advisable to lower the *min_template_size* (e.g., to 4). KS4 considers 5 different spatial ranges based on the *min_template_size* and checks for the best of 1, 2, 3, 4, and 5 times the *min_template_size* for spatial ranges during convolution of the simple templates. Because the default interchannel distance in EMUsort is 2 μm, changing this to 4 would cause the algorithm to check 4, 8, 12, 16, and 20 μm instead of the default 10, 20, 30, 40, and 50 μm, which would translate to focusing on roughly 2, 4, 6, 8, and 12 channels during convolution instead of roughly 5, 10, 15, 20, and 25 channels. We found better performance in sorting with the rat datasets when forcing the algorithm to use the wider spatial ranges (with *min_template_size* of 10) during the convolution of simple templates, likely related to the observation that most of the MUAPs in our biological data tend to span at least 5 channels.

### Methods for improved clustering of non-overlapping threshold crossings for simple template creation

As described in the Results section, selection of clean, representative waveforms for the simple templates is essential for all downstream stages of the KS4 spike sorting algorithm. In EMUsort we implemented changes that improved the robustness across our datasets in properly identifying unique MUAP waveforms for use as simple templates. The below 2 changes resulted in a larger initial set of non-overlapping threshold crossings, which improved the robustness of the outlier detection with HDBSCAN and subsequent K-means clustering accuracy (Campello et al., 2013, 2015).

First, we enabled the ability to use multiple single channel thresholds, as described in the Results, for greater detection of non-overlapping threshold crossings (i.e. those without neighboring threshold crossings within +/- 1 ms) even within regions of high motor unit activation, such as forceful behaviors. For example, if the user were to set a single threshold for *Th_single_ch* (e.g. [3] or [9], see Table 5), it is possible to miss larger MUAPs with the low threshold or to miss smaller MUAPs with the larger threshold (see Figure 4). The thresholds are in units of variance after channel whitening, so they are comparable to standard deviations. For the initial set of non-overlapping threshold crossings, a user should aim to find at least several hundred, generally getting better waveform initializations as the number increases. Adding more thresholds can increase the number of threshold crossings found, as can including lower thresholds. However, in our experience, using thresholds that are too low can increase noise in the sampling, including small noise fluctuations as crossings. The HDBSCAN removal can usually handle most of the cleanup through rejection of inconsistent waveforms, but it is generally recommended to avoid setting *Th_single_ch* below 3 standard deviations. It is also important to include higher thresholds, sometimes up to 12 standard deviations or more if the dataset of interest has a high dynamic range of MUAP amplitudes (e.g. largest spikes are more than ∼10 times bigger than smallest spikes) or if there are large and infrequently recruited MUAPs. When setting multiple thresholds, it is generally beneficial to separate the distance between threshold values by 2 or 3 standard deviations in order to avoid redundant threshold crossings and facilitate larger separations between clusters prior to creation of the simple templates. When adjusting this parameter, a user can inspect the cluster designations in Phy GUI and check the TraceView to ensure that no consistent MUAPs are failing to be classified into a cluster. If a small amplitude MUAP is being missed, it may be advisable to add a lower threshold for *Th_single_ch*. However, if some large amplitude MUAPs are missing, try a larger threshold. This parameter can also be explored with the parameter sweep function, followed by the selection procedure described in “Producing Composite Scores for Agnostic Estimation of Sort Quality” to guide parameter choices based on higher scores. Overall, this change facilitates producing a larger set of non-overlapping threshold crossings, by aggregation across thresholds, without including duplicate threshold crossing times.

Second, we greatly increased the number of batches to be included in the initialization stage. The default setting for *nskip* (see Table 5) in KS4 for subsampling batches is 25 (i.e., take all non-overlapping threshold crossings only from every 25th batch), but with MUAP datasets we found this to be too impoverished to effectively find a set of simple templates that spanned the entire range of spike waveform shapes. So, EMUsort uses every other batch (*nskip* set as 2 instead of 25). From our tests, we found that this change enabled HDBSCAN to perform more effectively in identifying true outliers by receiving input data with a much higher dynamic range of densities to operate on.

### Kilosort4 spike sorting details

#### Extraction of non-overlapping spikes for use as initial ‘simple templates’

The first stage of KS4 is to initialize the ‘simple templates’ (also referred to as universal templates), which have distinct waveform shapes for each candidate unit (number is determined by the *n_templates* parameter). They are expected to span the full range of waveform shapes present in the dataset so that no spikes are missed during convolutional template matching. Each of these simple templates is composed of a single waveform shape and a Gaussian spatial decay parameter that controls the amplitudes across different channels. The single waveform shape for each unit is duplicated for each channel, with the highest amplitude copy of the same shape being applied to a central best matching channel, with the same shape applied with descending amplitudes in all directions across the nearest channels according to a Gaussian distribution. Once these simple templates are learned, they will be convolved with the data to produce an initial set of spike times for each candidate unit. Simple templates are produced using the following approach.

After preprocessing (300 Hz high-pass filtering and whitening of channel amplitudes), a spike detection stage takes threshold crossings on single channels to determine spike times for waveform extraction, with the constraint that the spike waveforms must be time-separated from other spikes to be included in the set for initializing the simple templates. By default, this is a +/- 1 ms and +/- 6 channel region of isolation around each candidate spike. The threshold (*Th_universal*, see Table 5) is taken across each channel after channel whitening to variance 1. Therefore, *Th_universal* is in units of channel variance, so a single value is used for all data channels. This step produces peak locations for all non-overlapping threshold crossings in the recording. KS4 then extracts the voltage waveforms (2 ms by default, controlled by *nt*, see Table 5) at all non-overlapping peak locations.

#### Kilosort represents spike features using principal components from the non-overlapping spikes

Once the set of non-overlapping single-channel waveforms is extracted, it is used for two different purposes. First, the non-overlapping waveforms are used to define a set of PCs to be used as the temporal components (*W*) for all remaining spike feature extraction operations. Second, the set of waveforms is clustered by K-means and the waveform shape at each cluster center is taken to define the initial set of simple template shapes. Each simple template has a unique waveform shape centered on single best matching channels with decaying duplicated shapes in all directions on the channel map surrounding the best channel. These simple templates are used in the first data convolution to produce a set of initial spike times for each template by thresholding for high similarity peaks during convolution (*Th_universal*, see Table 5). These spike times are then used for multichannel feature extraction for each cluster by computing the PC loadings for small windows of data at each spike time using the same set of original PC components, applied to each channel. These multichannel PC loadings efficiently represent unique shapes for each of the clusters found so far, but they need to be differentiated and refined further to reduce the misclassifications that are likely present during this cluster initialization.

#### Kilosort classifies spike features with advanced clustering methods and spatiotemporal constraints

To properly split and merge the clusters, several measures are used to validate characteristics of normal spiking statistics in both timing and waveform shape. The initial multichannel PC loadings found by the previous stage are then clustered by KS4’s graph-based clustering and merging tree strategy as described previously (Pachitariu et al., 2024), and EMUsort introduces no new changes to this procedure. This cluster refinement process performs splits and merges of the clusters in PC space to prevent the presence of bimodality along the regression axis between PC loadings of the two clusters, which would indicate the presence of two distinct waveform shapes within a given cluster. After this process, the newly refined spike times are used to compute new multichannel PC loadings for each spike and all clusters. These loadings for each cluster are then used in the last convolutional template matching stage, where cleaner convolution outputs are extracted by progressively subtracting the best template matches, starting with the clusters that explain the most data variance. These subtractions leave behind a background of unaccounted for noise. This noise is later subtracted prior to computing the final PC loadings of the spikes in each cluster, which improves the accuracy of the shapes for the final cluster refinement stage. For each cluster, multichannel PC loadings or channel weights (*U*) are represented by the output of the convolution of each temporal component with each channel across all spike times of each cluster to define the magnitudes for the templates that correspond to each putative unit. Together, these low dimensional temporal components (PC components of the original non-overlapping waveforms) and channel weights (multichannel PC loadings) for each unit efficiently represent all spike shapes in the dataset. To perform the template matching, a 1D convolution is computed between each data channel and each temporal component along the time dimension, which results in a 3D output matrix (*num_chans*, *n_pcs*, *batch_size*). Immediately after, a linear summation is performed along several dimensions simultaneously (using the *einsum* implementation in Pytorch) to combine the 3D output matrix with the 3D matrix containing the channel weights (*n_units*, *n_pcs*, *num_chans*) to produce a 2D matrix (*n_units*, *batch_size*) which is the output of all combinations of template convolutions with all channels, to be thresholded for similarity by another variance threshold (*Th_learned*, see Table 5), producing the final set of spike times for each of these learned clusters corresponding to predicted units. There is another final stage of refinement that uses time and shape criteria to perform splits and merges of the clusters. KS4 computes cross-correlograms between spike times across clusters to minimize refractory period violations within each cluster. It also refines the spike shapes during merging by using the bimodality criteria on the PC loadings to decide the splitting again. With KS4, the clusters produced in the final output are generally overestimated so that a user typically receives more clusters in the output than the true number of underlying units. The user is expected to apply supervised selection of the best clusters that have the fewest numbers of errors such as refractory period violations (e.g., more than one spike occurring within 2 ms) or non-Gaussian spike amplitude distributions within a putative unit cluster.

## Acknowledgements

This work was supported by NIH grants R01NS109237 and U24NS126936 (to SJS), the Simons Foundation as part of the Simons-Emory International Consortium on Motor Control, the Emory School of Medicine through a Synergy II Nexus award, the Azrieli Foundation, Banting Research Foundation, Canada First Research Excellence Fund, Canada Research Chairs, Canadian Institutes of Health Research (PJT-175010), Natural Sciences and Engineering Research Council of Canada (RGPIN-2022-04421), and Vector Institute. We thank Mattia Rigotti and Yuyan (Margo) Shen for assistance in collecting rat datasets, Amanda Jacob for logistics, planning, and lab management necessary to be able to perform the rat surgeries for data collection, and Colleen Spellen for administrative support. We also thank the developers of MUedit for correspondence and providing feedback during the data analyses to produce the sorter comparisons.

## Additional Information

## Ethics

All experimental procedures described in this study were approved by the appropriate Institutional Animal Care and Use Committee at Emory University and Western University.

## Additional Files

## Supplementary Tables

**Table 1 - table supplement 1:** Descriptive statistics for distributions of cross-unit mean accuracy scores for each sort run with the rat simulated dataset, using the definition of accuracy as computed in (Pachitariu et al., 2024). Values in brackets were computed using only MUAPs that overlapped within half the template width of another MUAP.

| Descriptive Statistics for Distributions of Cross-Unit Mean Accuracy Scores |  |  |  |  |
| --- | --- | --- | --- | --- |
| Sorter | Run Count | Median (%)<br>all spikes [overlaps] | Mean (%)<br>all spikes [overlaps] | Standard Deviation (%)<br>all spikes [overlaps] |
| EMUsort | 25 | 94.8 [84.1] | 91.9 [79.8] | 6.2 [8.9] |
| Kilosort4 | 25 | 81.2 [51.1] | 76.4 [48.5] | 12.3 [13.4] |
| MUedit | 4 | 41.3 [16.3] | 45.0 [19.3] | 14.2 [15.7] |

**Table 1 - table supplement 2:** Descriptive statistics for distributions of cross-unit mean error rates for each sort run with the rat simulated dataset. This is computed by taking the accuracy scores from Table 1 - table supplement 1 for each sort run and subtracting them into 1. Values in brackets were computed using only MUAPs that overlapped within half the template width of another MUAP.

| Descriptive Statistics for Distributions of Cross-Unit Mean Error Rates |  |  |  |  |
| --- | --- | --- | --- | --- |
| Sorter | Run Count | Median (%)<br>all spikes [overlaps] | Mean (%)<br>all spikes [overlaps] | Standard Deviation (%)<br>all spikes [overlaps] |
| EMUsort | 25 | 5.2 [15.9] | 8.1 [20.2] | 6.2 [8.9] |
| Kilosort4 | 25 | 18.8 [48.9] | 23.6 [51.5] | 12.3 [13.4] |
| MUedit | 4 | 58.7 [83.7] | 55.0 [80.7] | 14.2 [15.7] |

**Table 1 - table supplement 3:** Per-unit performance statistics from the best EMUsort run, computed with all spikes from the rat simulated dataset.

| Unit | True Count | Sorter Count | Precision | Recall | Accuracy |
| --- | --- | --- | --- | --- | --- |
| 1 | 21704 | 20933 | 99.8% | 96.2% | 96.0% |
| 2 | 16484 | 15064 | 99.3% | 90.7% | 90.1% |
| 3 | 12529 | 12126 | 100.0% | 96.8% | 96.8% |
| 4 | 11041 | 10929 | 99.9% | 98.9% | 98.8% |
| 5 | 8438 | 8401 | 98.6% | 98.2% | 96.8% |
| 6 | 6052 | 5999 | 100.0% | 99.1% | 99.1% |
| 7 | 4190 | 4188 | 99.3% | 99.2% | 98.5% |
| 8 | 4143 | 4151 | 99.8% | 100.0% | 99.8% |
| 9 | 3178 | 3133 | 99.2% | 97.8% | 97.0% |
| 10 | 1330 | 1333 | 99.4% | 99.6% | 99.0% |
|  |  | **Median** | 99.6% | 98.5% | 97.7% |
|  |  | **Mean** | 99.5% | 97.7% | 97.2% |
|  |  | **Standard Deviation** | 0.5% | 2.7% | 2.8% |

**Table 1 - table supplement 4:** Per-unit performance statistics from the best sort run with EMUsort, computed only with overlapping spikes from the rat simulated dataset.

| **Unit** | **True Count** | **Sorter Count** | **Precision** | **Recall** | **Accuracy** |
| --- | --- | --- | --- | --- | --- |
| 1 | 8465 | 7150 | 96.7% | 81.7% | 79.5% |
| 2 | 7651 | 6195 | 95.7% | 77.5% | 74.8% |
| 3 | 6688 | 5941 | 97.9% | 87.0% | 85.4% |
| 4 | 6194 | 5907 | 97.5% | 93.0% | 90.8% |
| 5 | 5235 | 5118 | 95.6% | 93.4% | 89.5% |
| 6 | 4092 | 3958 | 98.4% | 95.2% | 93.7% |
| 7 | 3379 | 3382 | 96.1% | 96.2% | 92.6% |
| 8 | 3351 | 3362 | 96.6% | 96.9% | 93.7% |
| 9 | 2440 | 2341 | 95.0% | 91.1% | 86.9% |
| 10 | 1114 | 1108 | 97.8% | 97.3% | 95.3% |
|  |  | **Median** | 96.7% | 93.2% | 90.2% |
|  |  | **Mean** | 96.7% | 90.9% | 88.2% |
|  |  | **Standard Deviation** | 1.1% | 6.8% | 6.7% |

**Table 1 - table supplement 5:** Per-unit performance statistics from the best KS4 run, computed with all spikes from the rat simulated dataset.

| **Unit** | **True Count** | **Sorter Count** | **Precision** | **Recall** | **Accuracy** |
| --- | --- | --- | --- | --- | --- |
| 1 | 21704 | 21189 | 99.5% | 97.2% | 96.7% |
| 2 | 16484 | 15205 | 99.9% | 92.2% | 92.1% |
| 3 | 12529 | 12273 | 100.0% | 97.9% | 97.9% |
| 4 | 11041 | 10956 | 98.8% | 98.0% | 96.8% |
| 5 | 8438 | 6837 | 90.7% | 73.5% | 68.4% |
| 6 | 6052 | 5845 | 98.3% | 95.0% | 93.4% |
| 7 | 4190 | 4117 | 97.3% | 95.6% | 93.1% |
| 8 | 4143 | 4125 | 98.4% | 98.0% | 96.5% |
| 9 | 3178 | 3056 | 96.0% | 92.4% | 89.0% |
| 10 | 1330 | 1314 | 99.9% | 98.7% | 98.6% |
|  |  | **Median** | 98.6% | 96.4% | 95.0% |
|  |  | **Mean** | 97.9% | 93.8% | 92.3% |
|  |  | **Standard Deviation** | 2.8% | 7.5% | 8.9% |

**Table 1 - table supplement 6:** Per-unit performance statistics from the best sort run with KS4, computed only with overlapping spikes from the rat simulated dataset.

| Unit | True Count | Sorter Count | Precision | Recall | Accuracy |
| --- | --- | --- | --- | --- | --- |
| 1 | 8465 | 7352 | 87.4% | 75.9% | 68.4% |
| 2 | 7651 | 6212 | 92.0% | 74.7% | 70.2% |
| 3 | 6688 | 6114 | 92.1% | 84.2% | 78.5% |
| 4 | 6194 | 5816 | 90.4% | 84.8% | 77.8% |
| 5 | 5235 | 3761 | 75.7% | 54.4% | 46.3% |
| 6 | 4092 | 3691 | 89.5% | 80.8% | 73.8% |
| 7 | 3379 | 3220 | 93.6% | 89.2% | 84.1% |
| 8 | 3351 | 3243 | 93.6% | 90.6% | 85.4% |
| 9 | 2440 | 2260 | 85.8% | 79.5% | 70.3% |
| 10 | 1114 | 1068 | 92.6% | 88.8% | 82.9% |
|  |  | **Median** | 91.2% | 82.5% | 75.8% |
|  |  | **Mean** | 89.3% | 80.3% | 73.8% |
|  |  | **Standard Deviation** | 5.4% | 10.6% | 11.4% |

**Table 1 - table supplement 7:**
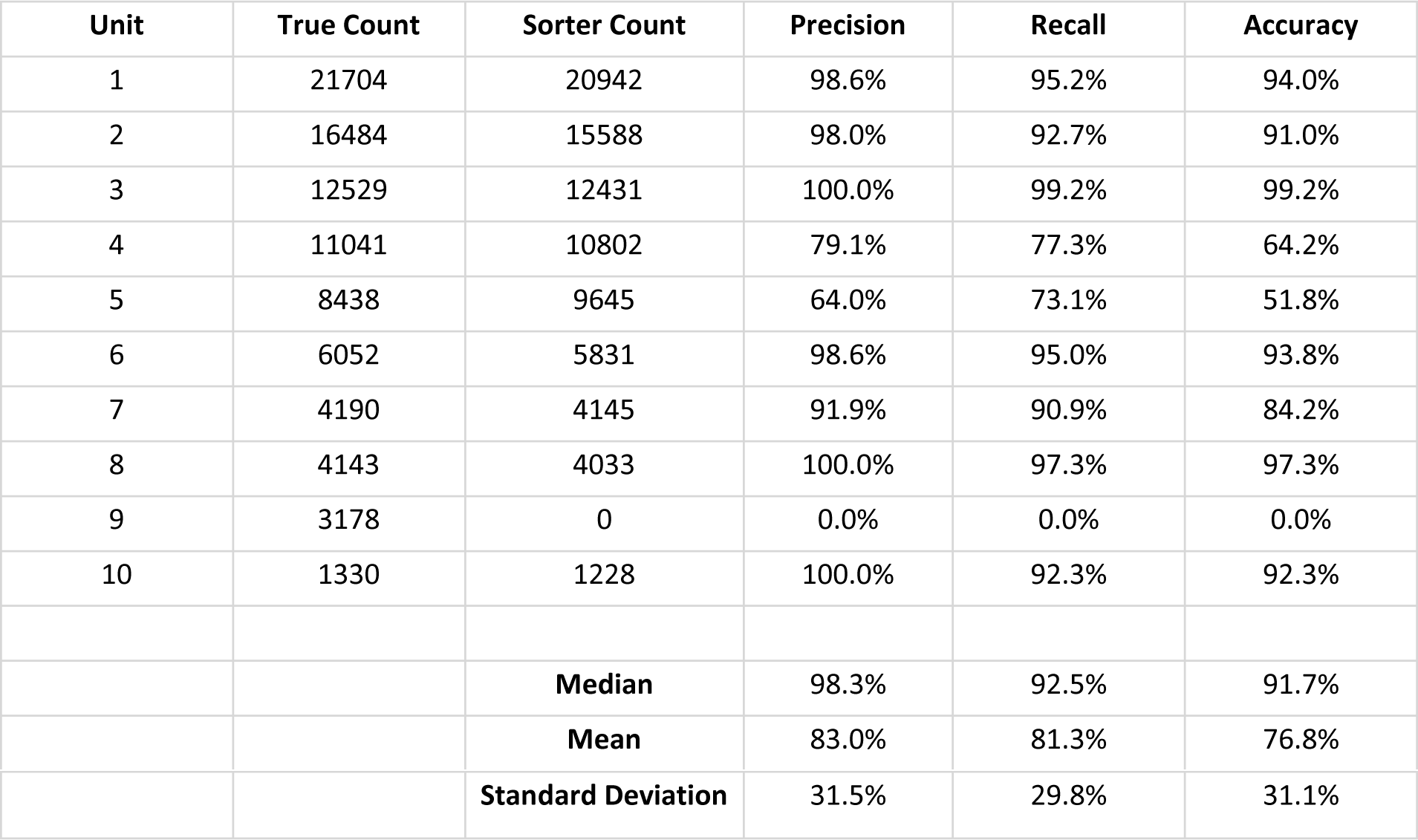
Per-unit performance statistics from the best MUedit run, computed with all spikes from the rat simulated dataset.

**Table 1 - table supplement 8:** Per-unit performance statistics from the best sort run with MUedit, computed only with overlapping spikes from the rat simulated dataset.

| Unit | True Count | Sorter Count | Precision | Recall | Accuracy |
| --- | --- | --- | --- | --- | --- |
| 1 | 8465 | 7277 | 85.1% | 73.2% | 64.9% |
| 2 | 7651 | 6787 | 85.8% | 76.1% | 67.6% |
| 3 | 6688 | 6645 | 86.3% | 85.7% | 75.4% |
| 4 | 6194 | 6371 | 53.7% | 55.2% | 37.4% |
| 5 | 5235 | 6788 | 47.3% | 61.4% | 36.5% |
| 6 | 4092 | 3787 | 82.0% | 75.9% | 65.0% |
| 7 | 3379 | 3317 | 83.6% | 82.1% | 70.7% |
| 8 | 3351 | 3755 | 83.7% | 93.8% | 79.3% |
| 9 | 2440 | 0 | 0.0% | 0.0% | 0.0% |
| 10 | 1114 | 964 | 94.4% | 81.7% | 77.9% |
|  |  | **Median** | 83.7% | 76.0% | 66.3% |
|  |  | **Mean** | 70.2% | 68.5% | 57.5% |
|  |  | **Standard Deviation** | 28.9% | 26.6% | 25.3% |

**Table 2 - table supplement 1:**
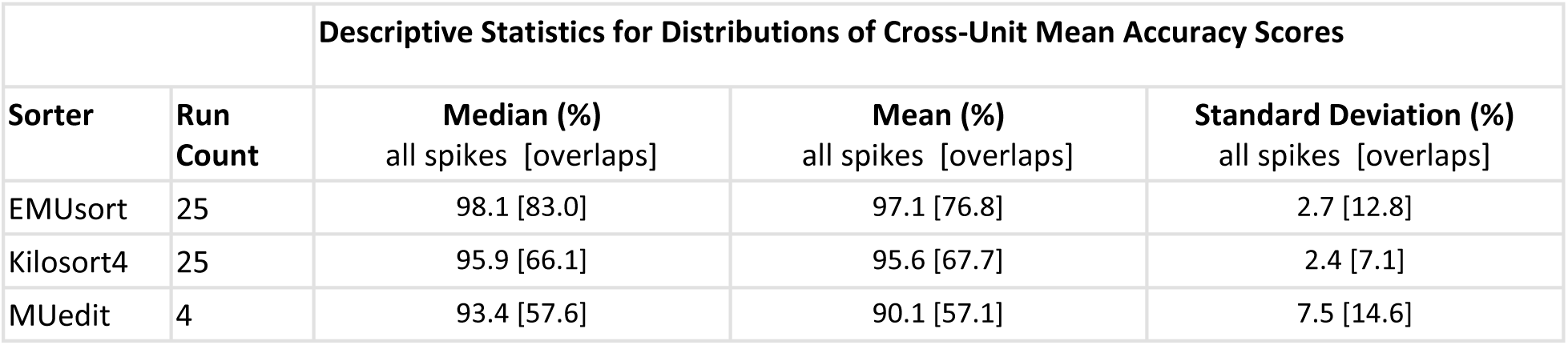
Descriptive statistics for distributions of cross-unit mean accuracy scores for each sort run with the monkey simulated dataset, using the definition of accuracy as computed in (Pachitariu et al., 2024). Values in brackets were computed using only MUAPs that overlapped within half the template width of another MUAP.

**Table 2 - table supplement 2:** Descriptive statistics for distributions of cross-unit mean error rates for each sort run with the monkey simulated dataset. This is computed by taking the accuracy scores from each sort run and subtracting them into 1. Values in brackets were computed using only MUAPs that overlapped within half the template width of another MUAP.

|  |  | **Descriptive Statistics for Distributions of Cross-Unit Mean Error Rates** |  |  |
| --- | --- | --- | --- | --- |
| Sorter | Run Count | **Median (%)**<br>all spikes [overlaps] | **Mean (%)**<br>all spikes [overlaps] | **Standard Deviation (%)**<br>all spikes [overlaps] |
| EMUsort | 25 | 1.9 [17.0] | 2.9 [23.2] | 2.7 [12.8] |
| Kilosort4 | 25 | 4.1 [33.9] | 4.4 [32.3] | 2.4 [7.1] |
| MUedit | 4 | 6.6 [42.4] | 9.9 [42.9] | 7.5 [14.6] |

**Table 2 - table supplement 3:** Per-unit performance statistics from the best EMUsort run, computed with all spikes from the monkey simulated dataset.

| Unit | True Count | Sorter Count | Precision | Recall | Accuracy |
| --- | --- | --- | --- | --- | --- |
| 1 | 12033 | 12038 | 99.9% | 99.9% | 99.8% |
| 2 | 6426 | 6421 | 99.9% | 99.8% | 99.7% |
| 3 | 3151 | 3151 | 100.0% | 100.0% | 100.0% |
| 4 | 1960 | 1960 | 100.0% | 100.0% | 100.0% |
| 5 | 645 | 645 | 100.0% | 100.0% | 100.0% |
|  |  | **Median** | 100.0% | 100.0% | 100.0% |
|  |  | **Mean** | 100.0% | 99.9% | 99.9% |
|  |  | **Standard Deviation** | 0.1% | 0.1% | 0.1% |

**Table 2 - table supplement 4:** Per-unit performance statistics from the best sort run with EMUsort, computed only with overlapping spikes from the monkey simulated dataset.

| Unit | True Count | Sorter Count | Precision | Recall | Accuracy |
| --- | --- | --- | --- | --- | --- |
| 1 | 1736 | 1754 | 93.3% | 94.2% | 88.2% |
| 2 | 1552 | 1559 | 92.7% | 93.1% | 86.7% |
| 3 | 1085 | 1080 | 96.2% | 95.8% | 92.3% |
| 4 | 780 | 774 | 97.2% | 96.4% | 93.8% |
| 5 | 290 | 295 | 96.6% | 98.3% | 95.0% |
|  |  | **Median** | 96.2% | 95.8% | 92.3% |
|  |  | **Mean** | 95.2% | 95.6% | 91.2% |
|  |  | **Standard Deviation** | 2.1% | 2.0% | 3.6% |

**Table 2 - table supplement 5:**
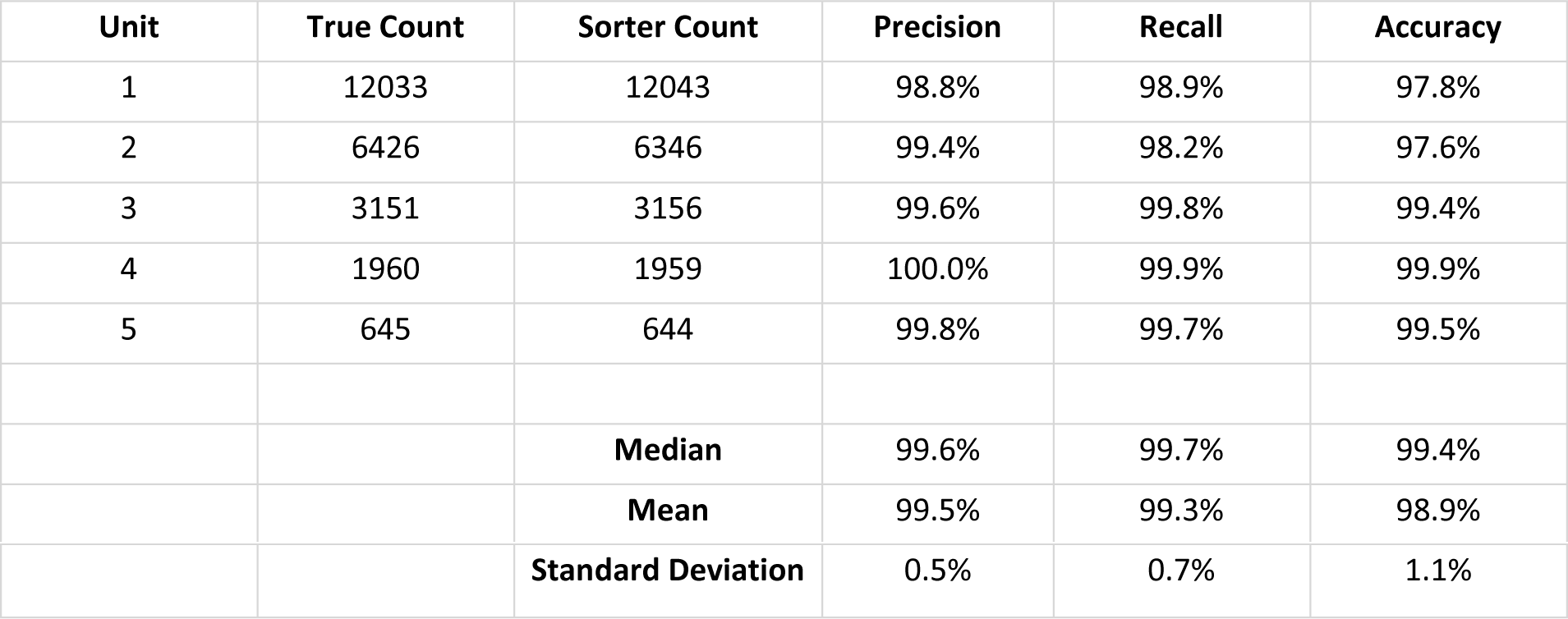
Per-unit performance statistics from the best KS4 run, computed with all spikes from the monkey simulated dataset.

| Unit | True Count | Sorter Count | Precision | Recall | Accuracy |
| --- | --- | --- | --- | --- | --- |
| 1 | 12033 | 12043 | 98.8% | 98.9% | 97.8% |
| 2 | 6426 | 6346 | 99.4% | 98.2% | 97.6% |
| 3 | 3151 | 3156 | 99.6% | 99.8% | 99.4% |
| 4 | 1960 | 1959 | 100.0% | 99.9% | 99.9% |
| 5 | 645 | 644 | 99.8% | 99.7% | 99.5% |
|  |  | **Median** | 99.6% | 99.7% | 99.4% |
|  |  | **Mean** | 99.5% | 99.3% | 98.9% |
|  |  | **Standard Deviation** | 0.5% | 0.7% | 1.1% |

**Table 2 - table supplement 6:**
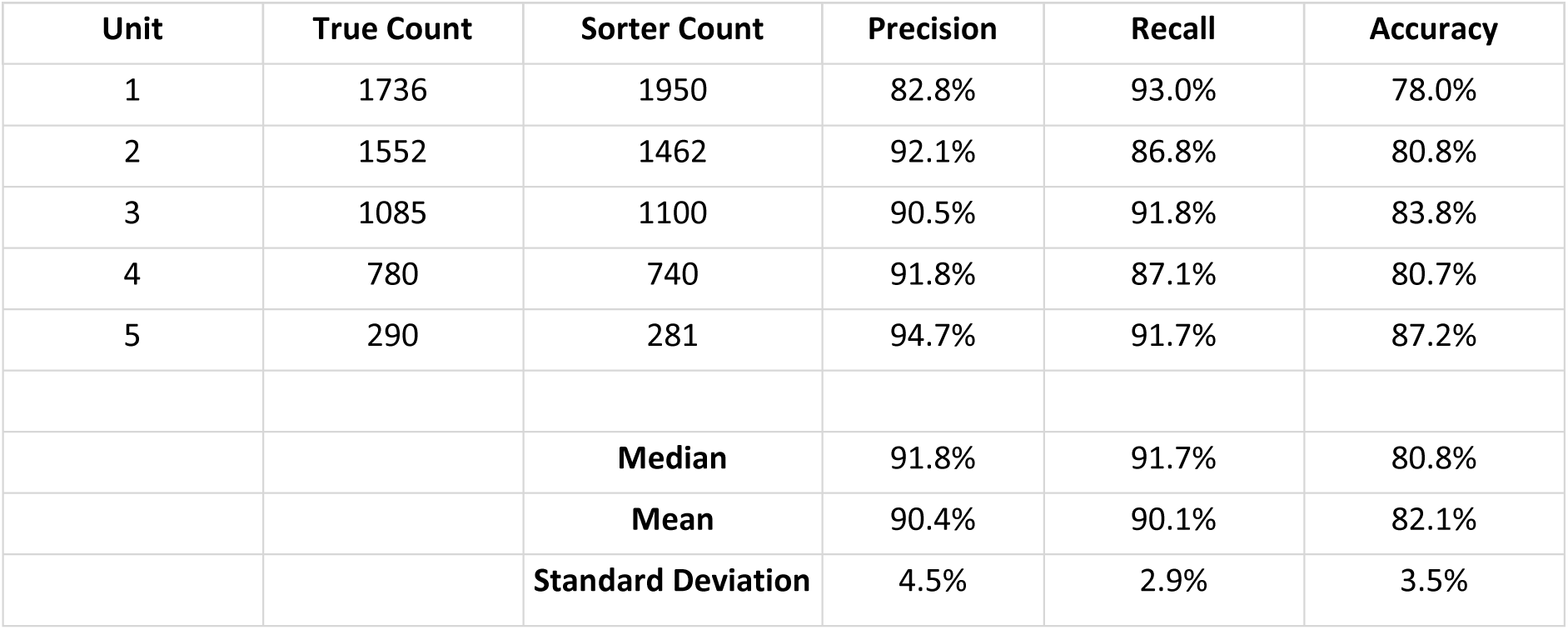
Per-unit performance statistics from the best sort run with KS4, computed only with overlapping spikes from the monkey simulated dataset.

| Unit | True Count | Sorter Count | Precision | Recall | Accuracy |
| --- | --- | --- | --- | --- | --- |
| 1 | 1736 | 1950 | 82.8% | 93.0% | 78.0% |
| 2 | 1552 | 1462 | 92.1% | 86.8% | 80.8% |
| 3 | 1085 | 1100 | 90.5% | 91.8% | 83.8% |
| 4 | 780 | 740 | 91.8% | 87.1% | 80.7% |
| 5 | 290 | 281 | 94.7% | 91.7% | 87.2% |
|  |  | **Median** | 91.8% | 91.7% | 80.8% |
|  |  | **Mean** | 90.4% | 90.1% | 82.1% |
|  |  | **Standard Deviation** | 4.5% | 2.9% | 3.5% |

**Table 2 - table supplement 7:** Per-unit performance statistics from the best MUedit run, computed with all spikes from the monkey simulated dataset.

| Unit | True Count | Sorter Count | Precision | Recall | Accuracy |
| --- | --- | --- | --- | --- | --- |
| 1 | 12033 | 11144 | 100.0% | 92.6% | 92.6% |
| 2 | 6426 | 6657 | 94.8% | 98.2% | 93.2% |
| 3 | 3151 | 2991 | 100.0% | 94.9% | 94.9% |
| 4 | 1960 | 1893 | 100.0% | 96.6% | 96.6% |
| 5 | 645 | 620 | 100.0% | 96.1% | 96.1% |
|  |  | **Median** | 100.0% | 96.1% | 94.9% |
|  |  | **Mean** | 99.0% | 95.7% | 94.7% |
|  |  | **Standard Deviation** | 2.3% | 2.1% | 1.8% |

**Table 2 - table supplement 8:** Per-unit performance statistics from the best sort run with MUedit, computed only with overlapping spikes from the monkey simulated dataset.

| Unit | True Count | Sorter Count | Precision | Recall | Accuracy |
| --- | --- | --- | --- | --- | --- |
| 1 | 1736 | 1555 | 97.6% | 87.4% | 85.6% |
| 2 | 1552 | 1747 | 77.8% | 87.6% | 70.1% |
| 3 | 1085 | 981 | 98.1% | 88.7% | 87.1% |
| 4 | 780 | 902 | 77.7% | 89.9% | 71.5% |
| 5 | 290 | 299 | 80.6% | 83.1% | 69.3% |
|  |  | **Median** | 80.6% | 87.6% | 71.5% |
|  |  | Mean | 86.4% | 87.3% | 76.7% |
|  |  | Standard Deviation | 10.5% | 2.6% | 8.9% |

## Supplementary Figures

**Supplementary Figure 1.**
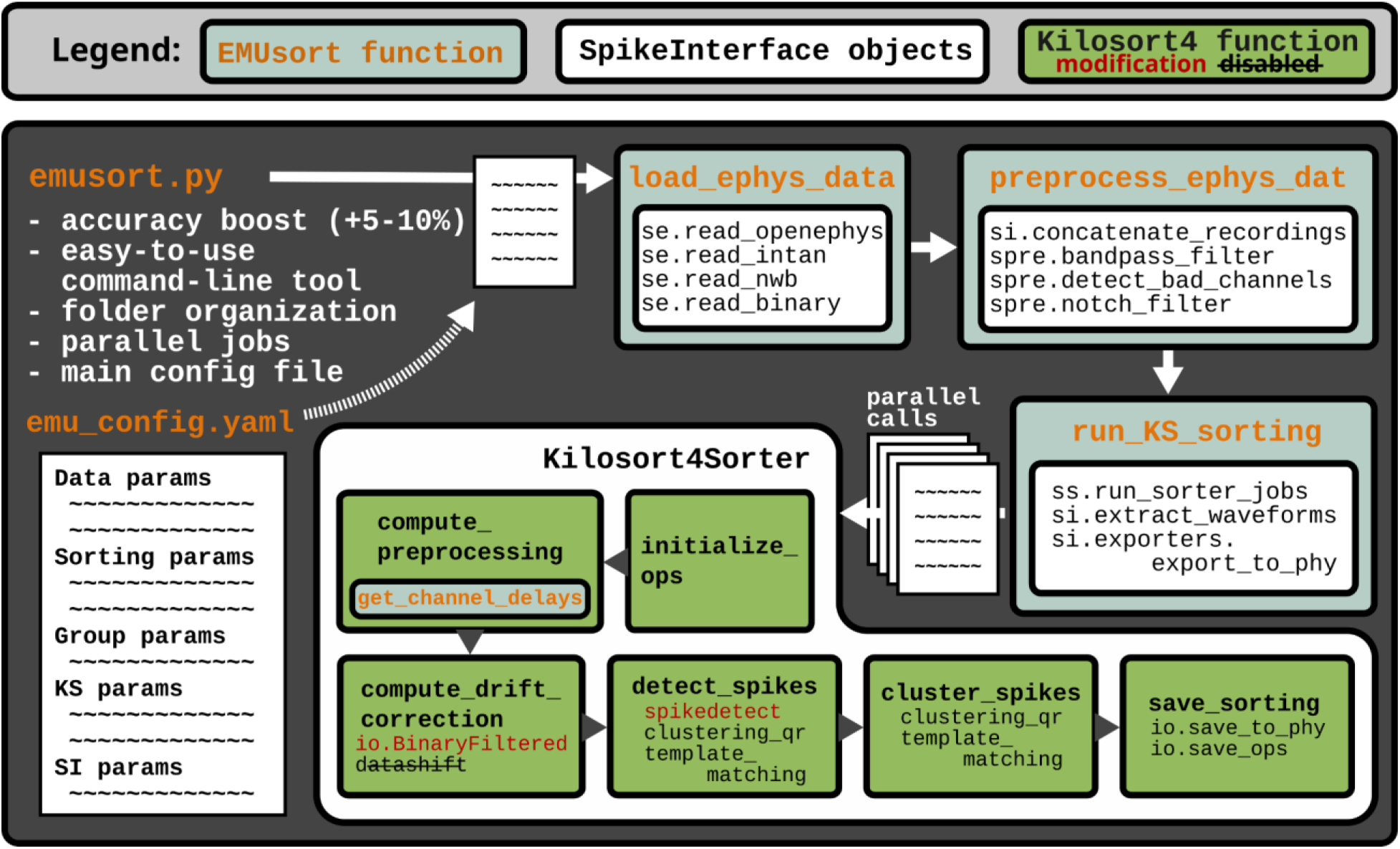
EMUsort leverages open-source libraries with a modular organization. The legend at the top indicates the color coding and symbolic meanings of the various boxes and fonts. The arrows indicate the order and hierarchy of function calls. The main script for the command line interface, “emusort.py”, utilizes a configuration file which stores all settings defining the input dataset properties, sorting parameters, settings defining separate groups of channels to use, and finally KS4 and SpikeInterface parameters. SpikeInterface methods are leveraged for loading, preprocessing, and starting KS4 processes in series or as parallel jobs with different settings. The modifications to KS4 source code, indicated by the legend, were implemented to remove channel delays and improve the initialization of templates.

**Supplementary Figure 2.**
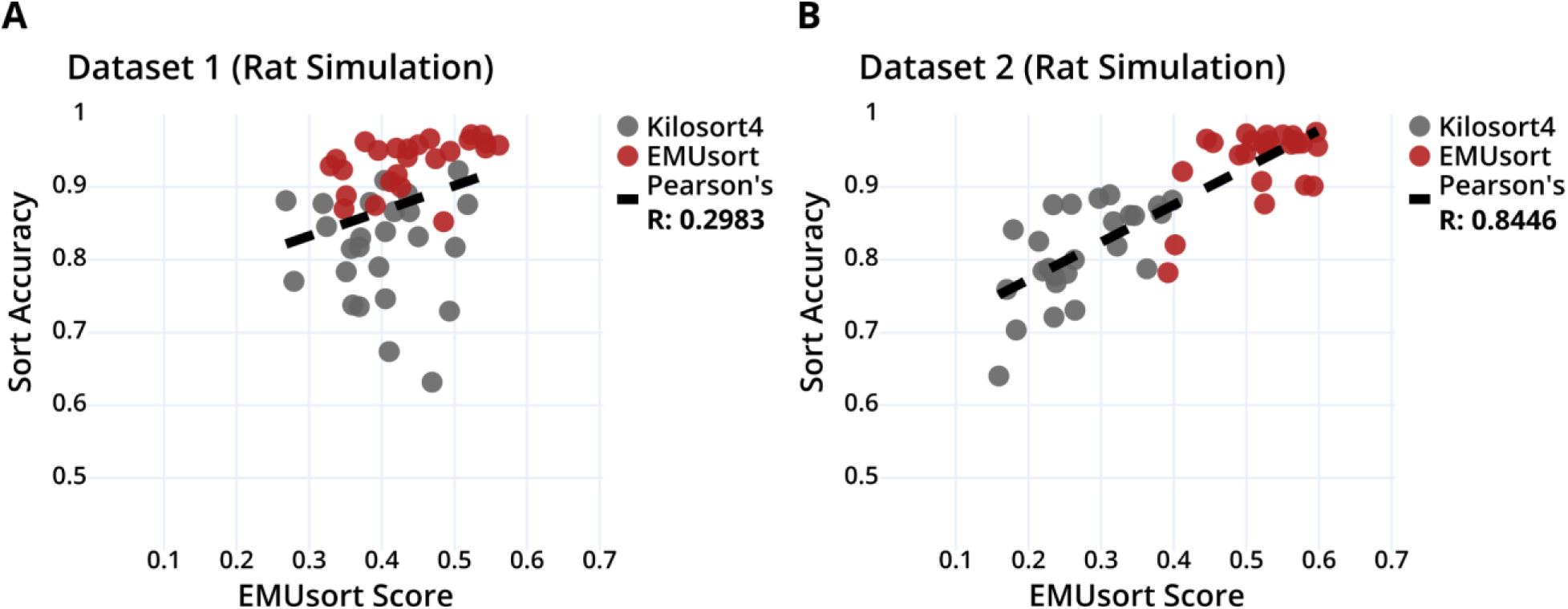
The composite scores produced by EMUsort for 2 of our simulated rat datasets can serve as a rough estimate of the computed ground truth accuracies (averaged across all units for each sort) with a weak positive correlation for Dataset 1 and a strong positive correlation for Dataset 2. Dataset 1 was used for the main results generated for Figure 6. Pearson’s R value is shown in the legend of each graph.

**Supplementary Figure 3.**
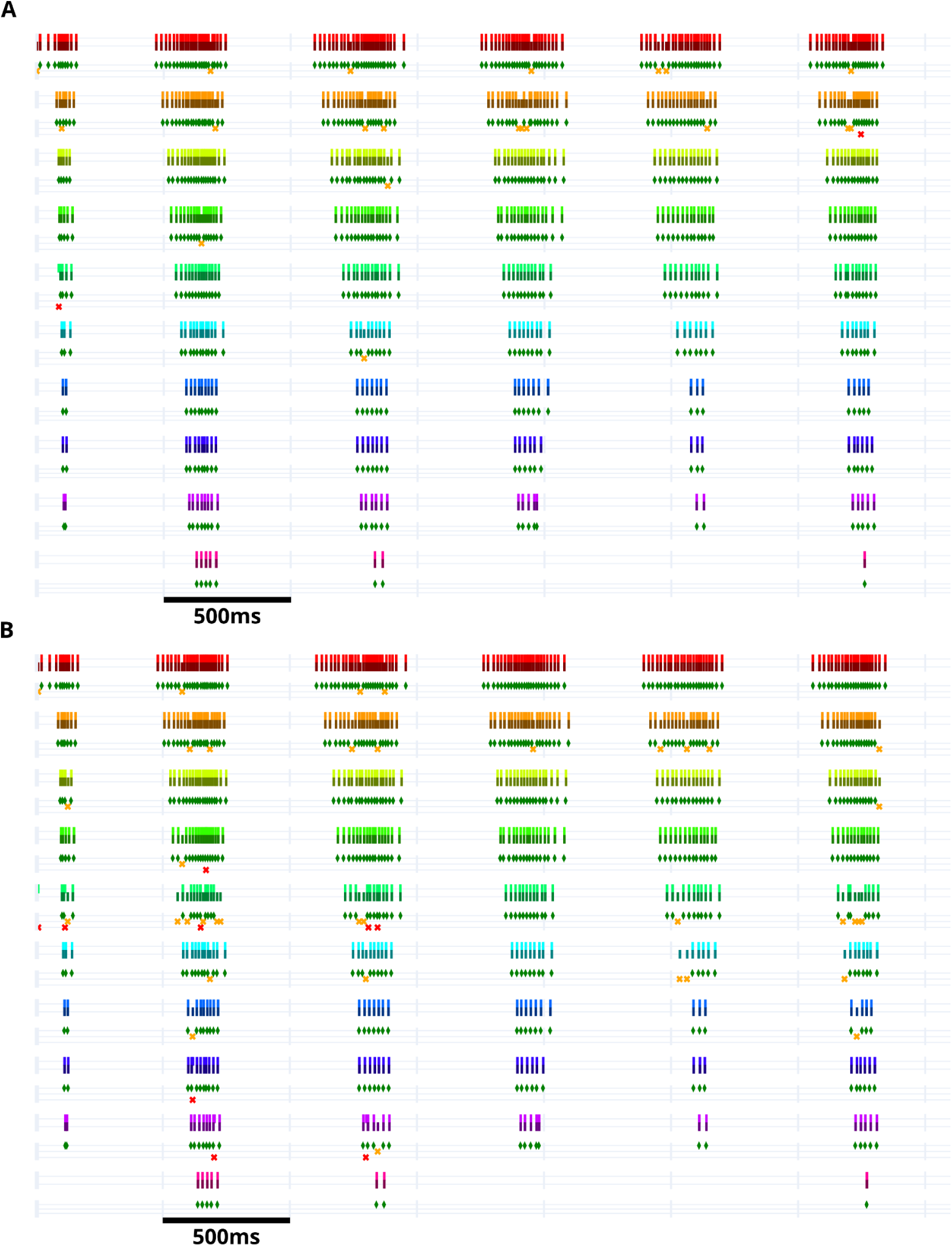
Example time stretch in the rat simulated dataset showing the ground truth versus identified spike times. Each row shows spike times with uniquely colored raster ticks for each motor unit. Darker ticks for each color show the ground truth spike times. Green diamonds mark correct times within +/- 1 ms. Yellow X’s mark false negatives. Red X’s mark false positives. (a) Event plot showing EMUsort evaluations. (b) Event plot showing KS4 evaluations.

## Data Availability

The EMUsort source code is available at https://github.com/snel-repo/EMUsort (also archived at (O’Connell & Michaels, 2025)). The code and instructions necessary to generate and access both the rat and monkey simulations are available at https://github.com/snel-repo/MUsim (also archived at (O’Connell, 2025)). All spreadsheet data used for computing (Tables 1–2), (Figure 6A-B), and (Figure 7A-B) are provided in the *EMUsort_paper_supplements* folder of the MUsim repository, with an explanation in the *README.md* file. The full original electromyography recordings and synchronized motion tracking data used to generate the rat simulation are stored as a single DANDI Archive (DANDI:001688/0.251231.0329; (O’Connell & Wang, 2025)).

